# Uracil-Driven ROS signals activate larval TrpA1-B neurons to prime the Drosophila adult gustatory response to bacterial signal

**DOI:** 10.64898/2025.12.22.695905

**Authors:** Romane Milleville, Gérard Manière, Martine Berthelot-Grosjean, Yaël Grosjean, Julien Royet, C. Léopold Kurz

## Abstract

Animals perceive their environment through their sensory systems why rely on neuronal circuits and proteins, but can be modulated by internal physiological conditions or exogenous factors. We have shown that the presence of certain pathogenic bacteria in the larval gut can alter the specificity of bacterial peptidoglycan recognition in the adults that emerge from these larvae. However, the nature of the bacterial signal and the host cells and molecules involved in receiving and transducing this signal remained unknown. We identify here uracil as the bacterial metabolite responsible for this priming and demonstrate that the effect is mediated by Duox-dependent production of reactive oxygen species in the larval gut. We show that specific expression of TrpA1 isoforms in a Gr66a-expressing neuron of the larval terminal organ is required for the adult response. Together, these findings reveal uracil as the bacterial cue and the Duox/ROS and TrpA1/Gr66a modules as key mediators linking larval gut microbial signals and host integration to subsequent sensory system modulation in the adult.

**Highlights:**

- Bacterial uracil primes larvae to drive adult aversion to peptidoglycan
- Duox-dependent ROS in larval enterocytes are required for sensory priming
- TrpA1 B/C/E isoforms in a larval Gr66a+ neuron mediate priming
- Uracil induces Duox/ROS signals enabling inter-larval activation of TrpA1-B+ neurons

## Introduction

In natural environments, animals inhabit diverse ecological niches that are colonized by bacteria, viruses, and fungi [1]. Throughout their development and into adulthood, animals interact with, and sometimes host, these microbial co-inhabitants. This close association can be beneficial, as certain microbial communities positively influence vital physiological functions such as fertility, lifespan, and growth[2], [3], [4]. However, some microbes can also negatively impact an animal’s health and internal balance [5]. The ability to detect and respond to such potential microbial threats is a fundamental and conserved innate mechanism essential for animal survival. To protect themselves and their offspring, animals have developed refined cellular and humoral innate immune responses [6]. Defensive responses launched after microbial detection can incur energetic or physiological costs [7], [8] and are not always completely effective [9], [10], [11]. Early detection of environmental dangers and preparation to confront these threats can complement the canonical immune responses against pathogens and serve as a critical initial defence. The nervous system’s ability to perceive microbial threats enables animals to adopt behaviours that mitigate the effects of infection on themselves and their progeny, at either the individual or collective level. Insects such as ants and bees, for example, employ social and behavioural immunity, like grooming, to protect against infection [12], [13], [14], [15], [16], [17], [18]. A variety of sensory systems— including smell [19], [20] and sight [21], [22]—are engaged in detecting biological threats in both vertebrates and invertebrates.

Research on Drosophila, has demonstrated that hygienic grooming behaviours can be triggered by fungal molecules or bacterial contact cues, through distinct sensory receptors and neural pathways [23], [24], [25]. Volatile compounds, such as geosmin produced by potentially pathogenic fungi, can be detected by insect olfactory receptors and act as repellents that affect food intake and egg laying [20]. Bacteria are characterized by cell wall structures such as peptidoglycan (PGN) or lipopolysaccharide, which serve as crucial ligands for receptors allowing eukaryotes to distinguish them from other organisms [26], [27], [28], [29]. Notably, these receptors are found on both immune cells and neurons [30], [31], [32].

Multiple studies have shown that bacterial PGN, a core cell wall component, mediates numerous interactions between bacteria and flies [24], [31], [33], [34], [35], [36], [37], [38]. Recognition of PGN by members of the PGRP family activates NF-kB pathways in immune-competent cells, leading to the production of immune effectors and regulators. Recent work has revealed that similar ligand/receptor interactions also maintain a molecular dialogue between bacteria and neurons in the fly’s central and peripheral nervous systems [31], [34], [37]. Hence PGN detection by adult taste neurons triggers immediate aversive behaviour in adult flies. Our recent findings identify two types of gustatory neurons—*ppk23+* and *Gr66a+*— as being essential for this response, though they serve distinct roles. Using time-specific neuronal inactivation and *in vivo* calcium imaging, we demonstrate that *ppk23+* neurons directly detect PGN in adult flies, whereas *Gr66a+* neurons must be active and possess the transient receptor potential TrpA1 channel during the larval stage to establish PGN sensitivity in adults. Moreover, adult flies that develop from larvae raised in germ-free (axenic) conditions lose the ability to respond to bacterial PGN [37]. Interestingly, reintroducing a single bacterial species, the pathobiont *Levilactobacillus brevis*, but not *Lactiplantibacillus plantarum*, into germ-free larvae—but not into adults—restores this response [37]. Therefore, the larval experience with bacteria influences adult sensory capabilities toward a microbial component. These findings which suggest a larval host-microorganism interaction and integration as well as a critical influence of both genetic and environmental factors during larval development in shaping the sensory capabilities of adult flies, raise two important questions. The first is how the bacterial information acquired by the larva is transmitted through metamorphosis to the future adult. The second, which is the subject of this study, concerns the nature of the signal provided by only some bacterial species and the mechanisms by which this or these signal (s) prime the larva to give birth to adults capable of sensing PGN through their gustatory neurons, a property absent in germ free animals.

## Results

### Mutant bacteria that do not release uracil are no longer able to prime larvae

We previously reported that wild-type flies from larvae reared in the presence of *Levilactobacillus brevis (L. brevis) are* attracted to a 1 mM sucrose solution [37]. However, this reflex is considerably attenuated when the sucrose solution contains bacterial peptidoglycan at a concentration of 200 µg/mL, suggesting that PGN has a bitter valence for adult flies (Figure and [37]). We have previously demonstrated that PGN derived from bacteria is perceived by *ppk23+* neurons located in the proboscis of adult flies and triggers aversion. We also showed that adults derived from larvae raised under axenic conditions lose this capacity while continuing to react to caffeine, an aversive molecule, demonstrating that their gustatory system remains functional. Furthermore, the induced cohabitation of axenic larvae with *L. brevis*, but not with *Lactiplantibacillus plantarum*, is sufficient to restore the aversive response to PGN in adults [37]. This suggests that certain specific bacterial species can prime larvae to give rise to PGN-sensitive adults (S1A). To identify the nature of the bacterial priming signal(s), we used a mono-association protocol in which axenic larvae are reared in the presence of the desired bacterial strains or compounds. Emerged adults are tested for their ability to respond to PGN using the proboscis extension reflex (PER) as a read out for the gustatory response (S1B). While larval cultivation with *L. brevis* primes the animals, neither heat-killed *L. brevis* nor the supernatant of a *L. brevis* solution were able to reproduce the priming effects of live *L. brevis* (Figure 1A). In order to search for a putative priming inducer, we looked for molecular characteristics present in bacterial species capable of priming but absent in others. While, as mentioned above, the opportunistic pathobionts *L. brevis* is capable of priming axenic larvae, *L. plantarum*, considered a symbiont for drosophila, is not[37]. As previous reports demonstrate that the nucleoside catabolism pathway controlling bacterial uracil is an essential trigger for the transition from commensal to pathogen[39], [40], we tested whether it could also be involved in the priming of axenic larvae. To this end, and for the remainder of the study, we used *Erwinia carotorova* (also known as *Pectobacterium carotovorum*,strain *Ecc^15^*) for which valuable mutants with impact on uracil metabolism had previously been generated[41], [42]. Unlike wild-type bacterial strains that were capable of larval priming, two mutants (*Ecc^15^*Δ4 and *Ecc^15^pyrE::Tn5*) whose ability to release uracil is impaired, lost this property (Figure 1B). This result, which suggests that uracil is involved in priming, was confirmed by experiments showing that simply adding purified uracil at 1mM to the medium is sufficient to trigger priming in larvae raised under axenic conditions (Figure 1B). Loss of priming in uracil-release mutants and sufficiency of exogenous uracil implicate uracil as a key determinant under our conditions.

**Fig. 1.**
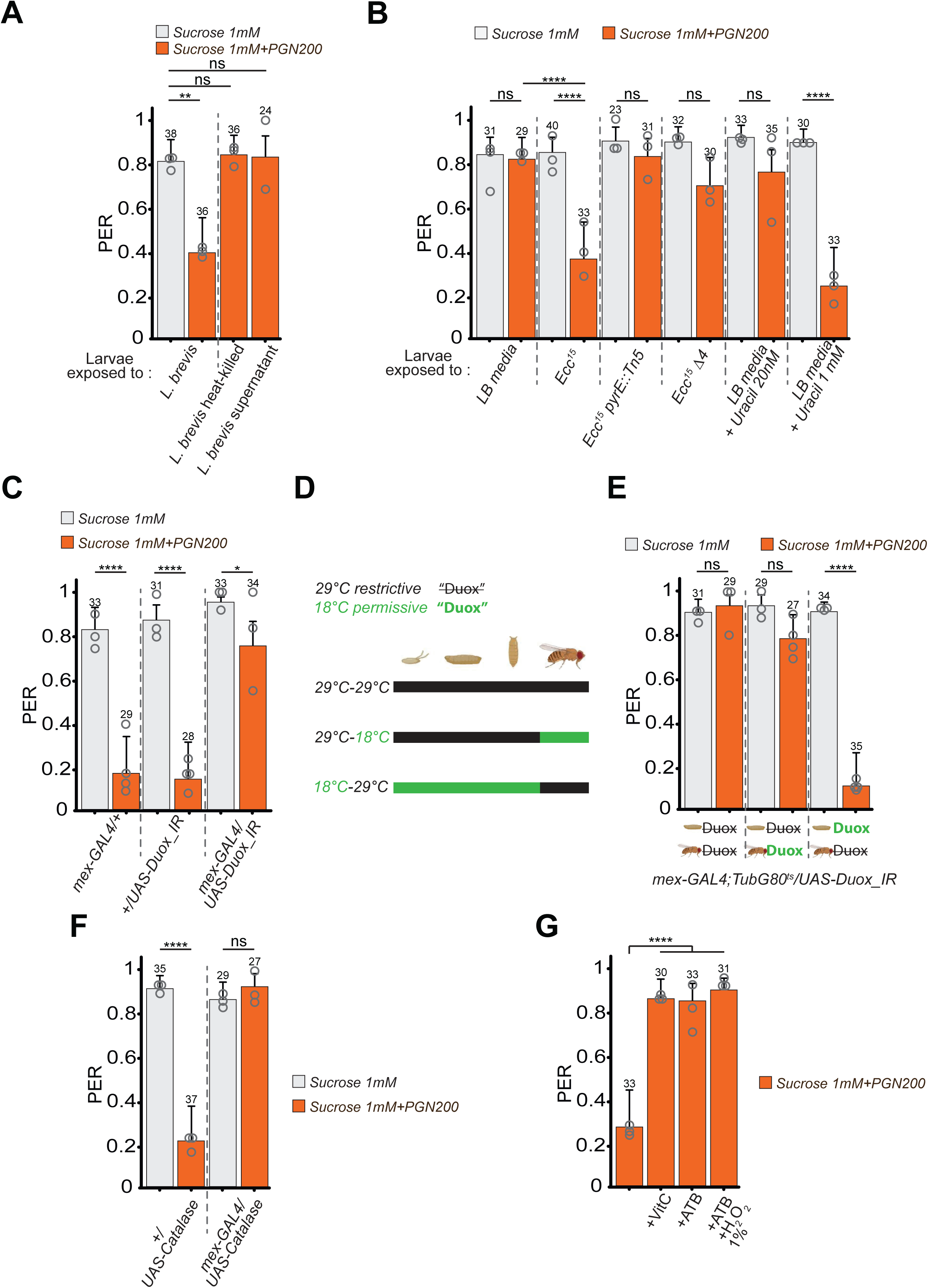
Larval exposure to uracil is required for adult PGN-mediated inhibition of PER. (**A**) Live *L. brevis* are necessary and sufficient to prime the larvae. PER index of w- flies to control solutions of sucrose and sucrose + PGN from *E. coli* K12 (PGN). Larvae from sterilized eggs were exposed to the different treatments and the resulting adults raised on antibiotics-containing media. (**B**) Uracil is necessary and sufficient to prime the larvae. PER index of w-flies to control solutions of sucrose and sucrose + PGN from *E. coli* K12 at 200µg/mL (PGN200). Larvae were monoassociated with bacterial strains, with LB media or with LB media containing uracil. The resulting adults were raised on antibiotics-containing media. **(C)** The Duox enzyme is necessary in enterocytes for the priming. PER index to solutions of sucrose and sucrose + PGN (PGN200) of *GAL4* (*mex*-*GAL4*/+) and *UAS (+/UAS-Duox_IR*) control flies as well as animals with the *Duox* transcript impaired via RNAi in enterocytes (*mex*-*GAL4*/*UAS-Duox_IR*). Larvae were raised on conventional media and the resulting adults raised on antibiotics-containing media. (**D**) Graphical representation of the life periods during which flies are shifted from 18***°***C to 29***°***C for a stage-dependent RNAi assay. The ubiquitously expressed *Tub-G80^ts^*, that inhibits the activity of GAL4, is temperature sensitive: the GAL4 inhibitor is active at 18***°***C (green) and inactivated at 29***°***C (black), its inactivation allows the expression of any *UAS*. In the case of *mex*-*GAL4*; *TubG80^ts^/UAS-Duox_IR*, at 18°C the *Duox* transcripts will not be impaired (**Duox**) while it will be at 29°C (). (**E**) Duox activity in enterocytes is necessary during the larval life for priming. PER index to solutions of sucrose and sucrose + PGN (PGN200) of *mex-GAL4; TubG80^ts^/UAS-Duox_IR* flies with the *Duox* transcript impaired all-life long, only during the larval stages or only during the adult stage. Larvae were raised on conventional media and the resulting adults raised on antibiotics-containing media. (**F**) Reducing the amount of ROS impairs the priming. PER index to solutions of sucrose and sucrose + PGN (PGN200) of *UAS (+/UAS-Catalase*) control flies as well as animals with the Catalase over-expressed in enterocytes (*mex-GAL4/UAS-Catalase*). Larvae were raised on conventional media and the resulting adults raised on antibiotics-containing media. (**G**) H_2_O_2_ supplementation to larvae is not sufficient to prime while vitamin C addition during the larval life prevents it. PER index to solutions of sucrose + PGN (PGN200) of *w^-^* flies. Larvae were raised on conventional media with or without vitamin C (0.2 mg/mL) or on antibiotics-containing media with or without H_2_O_2_ (1%). All the resulting adults were raised on antibiotics-containing media. Labellar PER was measured to 1 mM sucrose plus or minus PGN. The PER index is calculated as the percentage of flies tested that responded with a PER to the stimulation ± 95% confidence interval (CI). A PER value of 1 means that 100% of the tested flies extended their proboscis following contact with the mixture, a value of 0.2 means that 20% of the animals extended their proboscis. The number of tested flies (n) is indicated on top of each bar. For each condition, at least 3 groups with a minimum of 10 flies per group were used. Each independent group is represented as an open circle. ns indicates p>0.05, * indicates p<0.05, ** indicates p<0.01, *** indicates p<0.001, **** indicates p<0.0001 Fisher Exact Test. Further details can be found in the detailed lines, conditions and, statistics for the figure section.

### The production of reactive oxygen species by enterocytes is necessary for larval priming

Previous studies demonstrate that uracil acts as a microbe-derived factor that modulates intestinal immunity in *Drosophila*[39]. Uracil-dependent activation of Duox leads to the production in the intestinal tract of reactive oxygen species (ROS), which combat invading pathobionts but also induce damage to host tissues[42], [43], [44]. It has been shown that the inactivation of a single bacterial gene involved in uracil production is sufficient to induce a phenotypic change from a colitogenic bacterium to a commensal bacterium [45]. These results, which demonstrate the importance of Duox-dependent ROS production in mediating the effects of uracil, prompted us to test whether ROS were also involved in larval priming. To do this, we induced RNA interference-mediated reduction of *Duox* transcripts specifically in enterocytes using the *mex*-*GAL4* driver (Figure S2). While control adult strains showed the described aversion to PGN in the PER test, this was no longer the case for *mex-GAL4/UAS-Duox_IR* adult flies (Figure 1C). Taking advantage of the *GAL4*/*GAL80^ts^* binary system, we reduced ROS levels in either the larval or adult stage (Figure 1D and 1E). While reducing ROS levels in larval guts suppressed priming, this was not the case when the reduction was induced in the adult stage (Figure 1E). Priming in larvae was also suppressed by *GAL4*/*UAS*-mediated overexpression of catalase, an enzyme that degrades hydrogen peroxide, in the gut (Figure 1F). Similarly, the addition of the antioxidant vitamin C to the medium at a non-bactericidal concentration (Figure S3A) suppressed larval priming (Figure 1G). However, the addition of 1% H₂O₂ to axenic larvae was not sufficient to induce priming (Figure 1G and Figure S3B and C). Overall, these results support a model in which Duox-dependent ROS production in larval enterocytes is required for priming

### The B, C, and E isoforms of TrpA1, but not the A and D, are involved in larval priming

Our previous data demonstrate that expression of the TrpA1 ion channel in larval *Gr66a+* neurons is necessary for larval priming by bacteria [37]. Drosophila TrpA1, which shares conserved sensory functions with its mammalian ortholog and is activated by heat, ROS, UV light, and irritant chemicals (Figure 2A), is alternatively spliced to produce five isoforms (Figure 2B)[46], [47], [48], [49]. By generating and phenotypically analyzing knock-in flies expressing a single TrpA1 isoform and knock-out flies lacking selected TrpA1 isoforms, Gu and colleagues conclude that a given sensory stimulus preferentially activates a specific TrpA1 isoform *in vivo*[46]. Using this genetic toolkit, we sought to identify the number and nature of TrpA1 isoform (s) involved in larval priming to bacteria. Although these *TrpA1* mutants all responded similarly to sucrose, their PER to a sucrose+PGN solution differs. Among the *KI* lines, *B-KI*, *C-KI*, and *E-KI* flies reacted like the wild type. In contrast, flies expressing only isoform *A* or *D* showed a response similar to that of *TrpA1-KO* mutant flies, demonstrating that they lack the abxility to mediate larval priming. These results, which suggest that isoforms *B*, *C*, and *E*, but not *A* and *D*, are capable of priming, were confirmed by analysis of the different *KO* mutant flies (Figure 2C). Adults from lines in which at least one of the three competent isoforms was present (*E-KO*, *BC-KO*, and *AD-KO*) all responded to PGN like the controls. This result was consistent with the fact that *BCE-KO* flies, which lack all three competent isoforms (*B*, *C* and *E*), were the only flies whose ability to respond to PGN was impaired (Figure 2C). These results demonstrate that expression of any one of the three isoforms, *B, C*, or *E* is sufficient to ensure larval priming. The simplest explanation would be that B, C, and E are functionally redundant and share a functional property that is absent in isoforms A and D. Careful examination of the TrpA1 peptides suggests that this is not the case. Indeed, no protein domain that is specifically present in proteins TrpA1 B, C, and E but absent in A and D could be identified. An alternative hypothesis would be that the type of cells(s) in which the *TrpA1* isoforms are expressed is the important criterion to consider and that isoforms *B*, *C* and *E* are co-expressed in one or more cells that would be necessary to ensure TrpA1-dependent priming. It should be noted that these two hypotheses (the nature of the peptides or the domain of expression) are not mutually exclusive (see below).

**Fig. 2.**
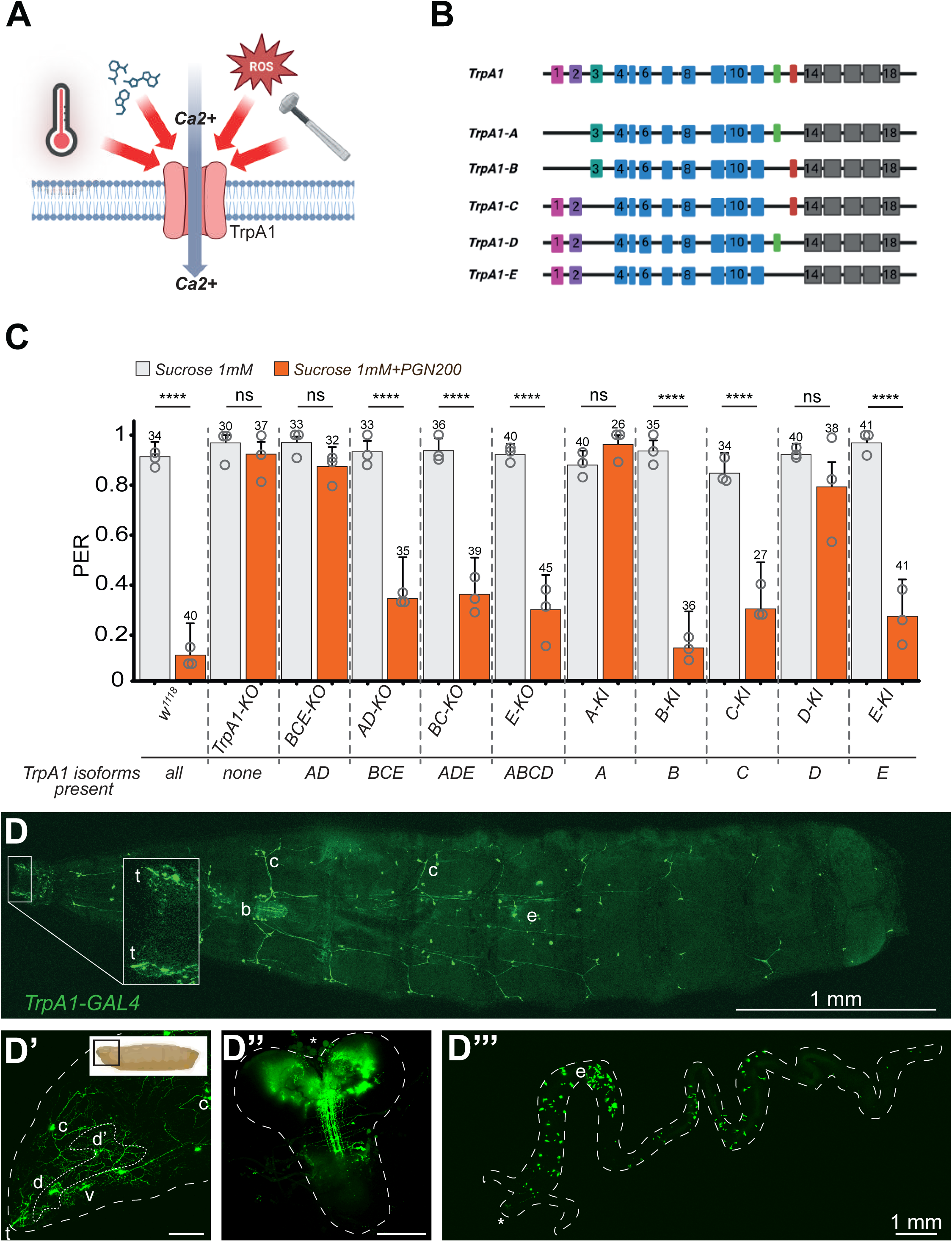
TrpA1 and specifically the B, C and E isoforms are involved in the priming. **(A**) Graphical representation of the TrpA1 channel responding to various stimuli through the entry of ions into the cell. Heat, specific chemicals, ROS and mechanical stimuli trigger the TrpA1 proteins participating to the generation of an action potential. **(B)** Graphical representation of the *TrpA1* gene (top) with colored boxes representing exons and colors related to functional domains of the translated protein. Five spliced variant isoforms have been reported so far, they all share exons 4-11 and exons 13-18. **(C)** *TrpA1-B*, *-C* and *-E* isoforms are involved in priming. PER index to solutions of sucrose and sucrose + PGN (PGN200) of control flies as well as *KO* and *KI* animals for the *TrpA1* isoforms. Depending on the genotype of the tested animal, the *TrpA1* isoforms that are present are listed under the x-axis. **(D-D’’’)** The *TrpA1-GAL4* driver (pan-isoforms) is widely expressed. Confocal images of larvae expressing *gfp* under the control of the pan-isoforms *TrpA1-GAL4* driver with a whole L3 larvae (**D**), the anterior part of a larvae with anteriormost on the left (**D’**), the larval brain (**D’’**) and the gut (**D’’’**). In (**D**), the larger rectangle is a magnification of the most anterior part of the animal housing two *gfp+* symmetrical neurons whose dendrites extend toward the external environment (t). (b) is for brain, (c) is for C4da neurons, (d) is for dorsal pharyngeal sensilla ganglion, (d’) is for dorsal pharyngeal organ ganglion, (v) is for ventral pharyngeal sensilla ganglion and (e) is for entero-endocrinal cells. In **D’** and **D’’**, the scale bar represents 100 µm. In **D’** and **D’’**, the asterisk indicates the anterior part of the organ. A schematic representation of the whole larva indicates the depicted area. The PER index is calculated as the percentage of flies tested that responded with a PER to the stimulation ± 95% confidence interval (CI). A PER value of 1 means that 100% of the tested flies extended their proboscis following contact with the mixture, a value of 0.2 means that 20% of the animals extended their proboscis. The number of tested flies (n) is indicated on top of each bar. For each condition, at least 3 groups with a minimum of 10 flies per group were used. Each independent group is represented as an open circle. ns indicates p>0.05, * indicates p<0.05, ** indicates p<0.01, *** indicates p<0.001, **** indicates p<0.0001 Fisher Exact Test. Further details can be found in the detailed lines, conditions and, statistics for the figure section.

### *TrpA1 B, C* and *E* isoforms are co expressed in some anterior neurons

As expected from a functionally pleiotropic protein, *TrpA1* is expressed in many cell types in both larvae and adults. In addition, Gu et al, have previously shown that each *TrpA1* isoform has a unique expression pattern and that multiple isoforms are often co-expressed in the same cells. To investigate whether the shared ability to prime of B, C, and E isoforms can be attributed to their co expression in some functionally important cells, we looked for cells that would express all 3 isoforms. For that, we crossed the *TrpA1*-isoform-*GAL4* flies with *UAS-gfp* and compared their expression domain. Interestingly, all 3 isoforms were expressed in a single and isolated anteriormost neuron probably belonging to one of the bitter neurons of the sensory Terminal Organ (TO) and in few neurons of the Dorsal and Ventral organs (Figure 3 A, B, C and figure 2D). Additionally, all three isoforms were also expressed in the horse shoe like shaped BLP larval brain neurons that have been shown to play a role in thermal nociception [51], while only *C* and *E* were expressed in the larval VNC (Figure 3 D, E and F). Finally, all three isoforms and the pan-isoforms were expressed in gut cells, which are likely to be entero-endocrinal cells (EEC) (Figure 2D and Figure 3G, H and I). However, while as reported earlier, *TrpA1-C* and *TrpA1-E* were expressed in C4da larval nociceptors neurons[46], it was not the case for *TrpA1-B* (Figure S4).

**Fig. 3.**
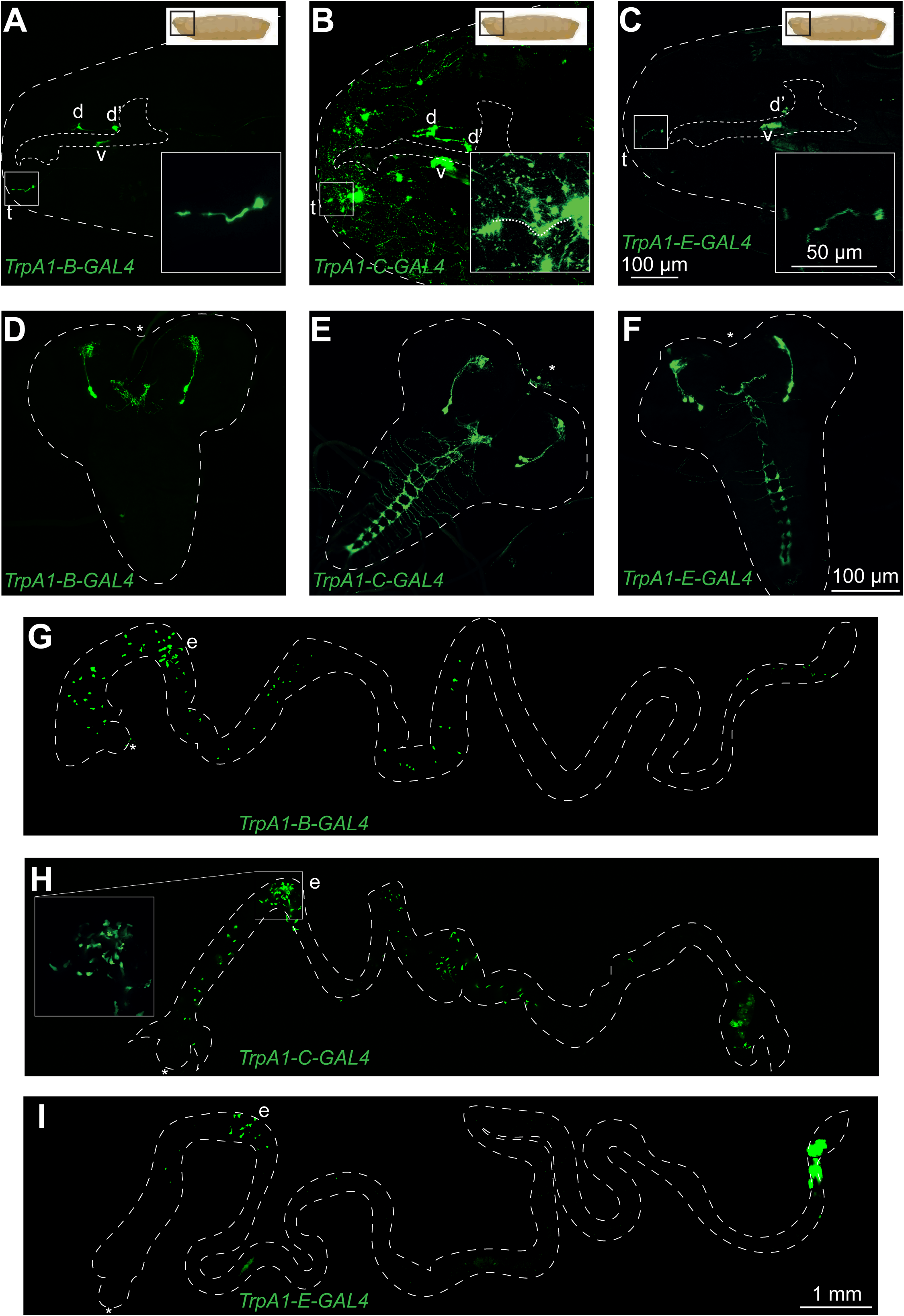
The expression pattern of the B isoform contains all the elements common to isoforms B, C and E. (**A-I**) Confocal images of larvae expressing *gfp* under the control of the *TrpA1-B-GAL4* driver (**A**, **D** and **G**), the *TrpA1-C-GAL4* driver (**B**, **E** and **H**) or the *TrpA1-E-GAL4* driver (**C**, **F** and **I**). Shown are the anterior parts (**A-C**), the brains (**D-F**) and the guts (**G-I**). In (**A-C**), the anterior is on the left and the larger square is a magnification of the most anterior part of the animal housing a neuron whose dendrite extends toward the external environment (t). In (**D-I**), the asterisk indicates the anterior part of the organ. (b) is for brain, (c) is for C4da neurons, (d) is for dorsal pharyngeal sensilla ganglion, (d’) is for dorsal pharyngeal organ ganglion, (v) is for ventral pharyngeal sensilla ganglion and (e) is for entero-endocrinal cells. In (**B**), strong *gfp* expression in C4da neurons makes it difficult to distinguish other transgene-expressing cells, but the neuron extending its dendrite toward the external environment is still detectable and outlined in the larger square. A schematic representation of the whole larva indicates the depicted area.

Since the *TrpA1-B KI* line has the most restricted expression pattern of the 3 reporter lines with no expression in the VNC and in the C4da larval nociceptors neurons, we further focused on this line. We first confirmed, via RNAi-mediated downregulation, that *TrpA1* inactivation in the larval but not the adult *TrpA1-B-GAL4* expression domain was sufficient to block priming (Figure 4A and 4B). This result, together with those previously reported, demonstrate that inactivating *TrpA1* in either *Gr66a-GAL4* [37] or in *TrpA1-B-GAL4* larval cells prevent larval priming. This suggests that *TrpA1-B-GAL4* and *Gr66a-GAL4* are co-expressed in one or more cells in which *TrpA1* expression would be required for priming. Consistently, using the binary system *GAL4*/*UAS* and *LexA/LexAop*, we showed that one of the three *Gr66a* positive neurons is *TrpA1-B-GAL4* (Figure 4C, C’ and C’’). We also excluded the gut part as being essential for priming since *Gr66a-GAL4* is not expressed in the gut ([37] and Figure S5). To further identify the neuron(s) and the TrpA1 isoform mediating priming, we try to rescue the pan isoform *TrpA1* mutant by providing different *TrpA1* isoforms in *Gr66a+* neurons. While the ability to prime that is lost in *TrpA1* mutant was fully rescued by expressing *TrpA1-B* in *Gr66a+* cells, this was not the case for *TrpA1-A* (Figure 4D and Figure S6). Altogether, these results have several implications. (i) they demonstrate that the larval neurons and not the enteroendocrine cells are required for priming (ii) they demonstrate that the expression of *TrpA1-B* in *Gr66a+* cells is sufficient for priming (iii) they strongly suggest that the *TrpA1-B*+/*Gr66a*+ anterior neuron is the cell in which *TrpA1-B* (or probably either *C* and *E*) need to be expressed to allow priming (iv) they demonstrate that all the *TrpA1* isoforms are not functionally substitutable as far as priming is concerned. All of these results were confirmed using another *GAL4* line (*Gr39a.b*), whose expression pattern in the larvae resembles that of *TrpA1-B* (Figure S7A) and which is not expressed in the EEC (Figure S7C). Co-expression confirmed that one of the *Gr39a.b-GAL4* TO neurons is *TrpA1-B* positive (Figure S7B). Moreover, impairing *TrpA1* expression in the *Gr39a.b*-*GAL4* cells that do not include EEC was sufficient to abrogate the adult PGN avoidance phenotype observed in controls (Figure 4E).

**Fig. 4.**
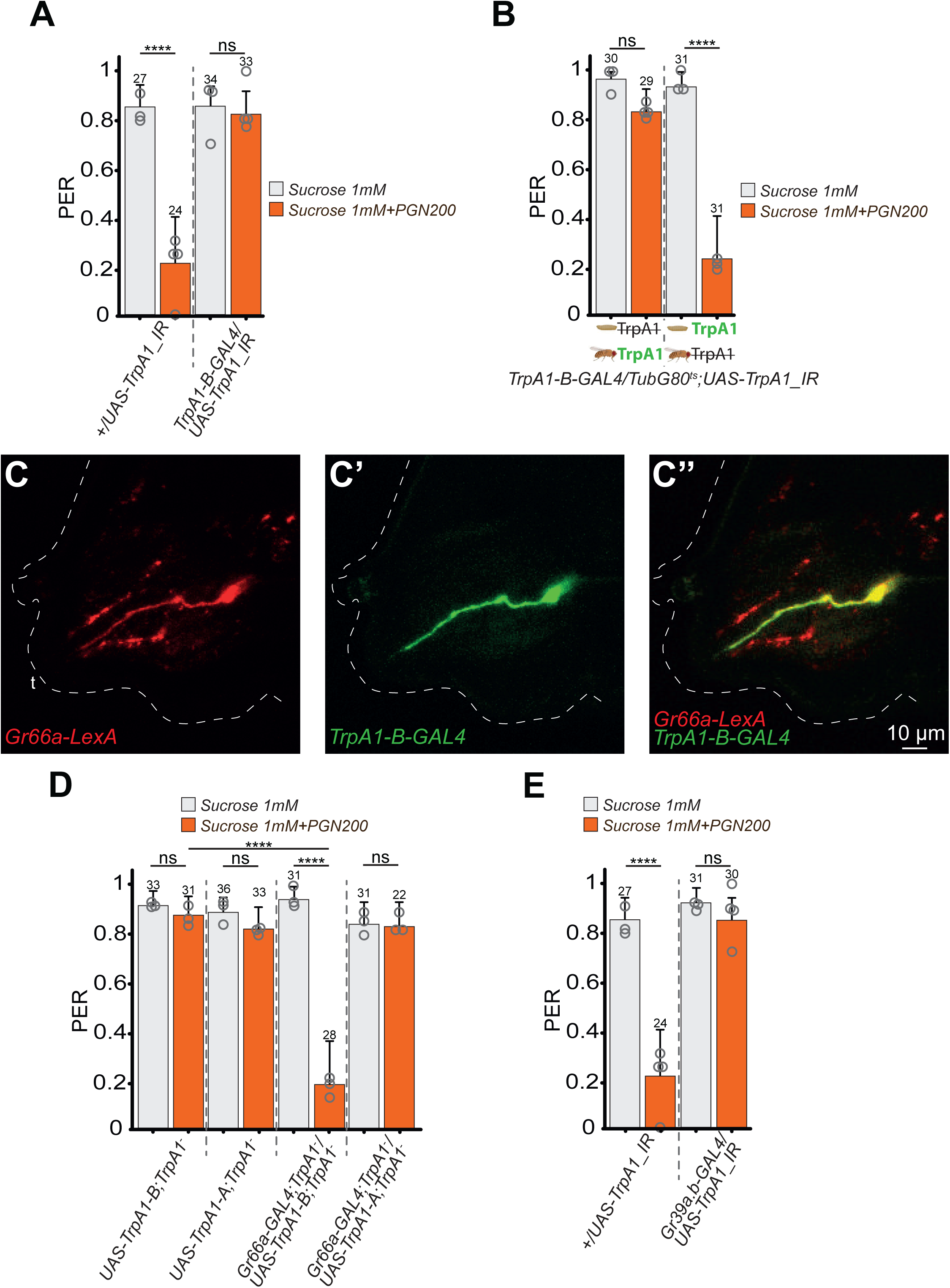
*TrpA1-B+* cells are *Gr66a+* as well as necessary for priming with the TrpA1-B isoform sufficient. **(A)** TrpA1 is necessary for priming in *TrpA1-B+* cells. PER index to solutions of sucrose and sucrose + PGN (PGN200) of control flies *(+/UAS-TrpA1_IR*) as well as animals with the *TrpA1* transcripts impaired via RNAi in *TrpA1-B+* cells (*TrpA1-B-GAL4/UAS-TrpA1_IR*). Larvae were raised on conventional media and the resulting adults raised on antibiotics-containing media. **(B)** *TrpA1* is necessary for priming in *TrpA1-B+* cells only during larval stage. PER index to solutions of sucrose and sucrose + PGN (PGN200) of *TrpA1-B-GAL4; TubG80^ts^/UAS-TrpA1_IR* animals with the *TrpA1* transcripts impaired via RNAi in *TrpA1-B+* cells only during the larval stages (left) or only during the adult stage (right). Larvae were raised on conventional media and the resulting adults raised on antibiotics-containing media. **(C-C’’)** Confocal images of the anterior part of a representative larva expressing *mCherry* under the control of the *Gr66a-LexA* driver as well as *gfp* under the control of *TrpA1-B-GAL4* driver. In **C**, dendrites from three *Gr66a+* neurons are detectable, but only one cellular body is clearly visible. **(D)** *TrpA1-B* isoform is sufficient for priming in *Gr66a+* cells while *TrpA1-A* is not. PER index to solutions of sucrose and sucrose + PGN (PGN200) of control flies impaired for priming (*UAS-TrpA1-B; TrpA1^-^* and *UAS-TrpA1-A; TrpA1^-^*) as well as animals with the *TrpA1* transcripts over-expressed only in *Gr66a+* cells in the *TrpA1^-^* mutant background (*Gr66a-GAL4; TrpA1^-^/ UAS-TrpA1-B; TrpA1^-^* and *Gr66a-GAL4; TrpA1^-^; UAS-TrpA1-A; TrpA1^-^*). Larvae were raised on conventional media and the resulting adults raised on antibiotics-containing media. (**E**) Impairing *TrpA1* expression in *Gr39a.b+* cells prevent priming. PER index to solutions of sucrose and sucrose + PGN (PGN200) of control flies (*+/UAS-TrpA1_IR*). Larvae were raised on conventional media and the resulting adults raised on antibiotics-containing media. The *+/UAS-TrpA1_IR* data from Figure 4E are those present on figure 4A. The PER index is calculated as the percentage of flies tested that responded with a PER to the stimulation ± 95% confidence interval (CI). A PER value of 1 means that 100% of the tested flies extended their proboscis following contact with the mixture, a value of 0.2 means that 20% of the animals extended their proboscis. The number of tested flies (n) is indicated on top of each bar. For each condition, at least 3 groups with a minimum of 10 flies per group were used. Each independent group is represented as an open circle. ns indicates p>0.05, * indicates p<0.05, ** indicates p<0.01, *** indicates p<0.001, **** indicates p<0.0001 Fisher Exact Test. Further details can be found in the detailed lines, conditions and, statistics for the figure section.

### The *TrpA1-B* positive TO neuron is activated by H_2_O_2_

After identifying, on the one hand, that *TrpA1-B* expression in the *TrpA1-B+/Gr66a+* anterior neuron is necessary for priming and, on the other hand, that ROS are also involved, we hypothesized that the latter could be the signal activating the former. To test this model, we used calcium imaging to measure the ability of *Gr66a+* and *TrpA1+* anterior neurons to respond to H₂O₂. When larvae expressing *Gr66a-GAL4*/*UAS-GCaMP* were stimulated with a 1% H₂O₂ solution, only one of the three anterior neurons responded strongly to the stimuli (Figure 5A-D). Interestingly, the neuron that responded to H₂O₂ was the only one to respond to quinine (Figure 5D and Figure S8). Using the same approach, we were able to show that the TO *TrpA1-B-GAL4* neuron was activatable by H₂O₂ and that this response was abrogated if *TrpA1* transcripts were cell specifically downregulated (Figure 5E-H). These results support that a single TO *TrpA1+/Gr66a+* neuron contributes to larval priming and is directly activated by ROS in a TrpA1-dependent manner.

**Fig. 5.**
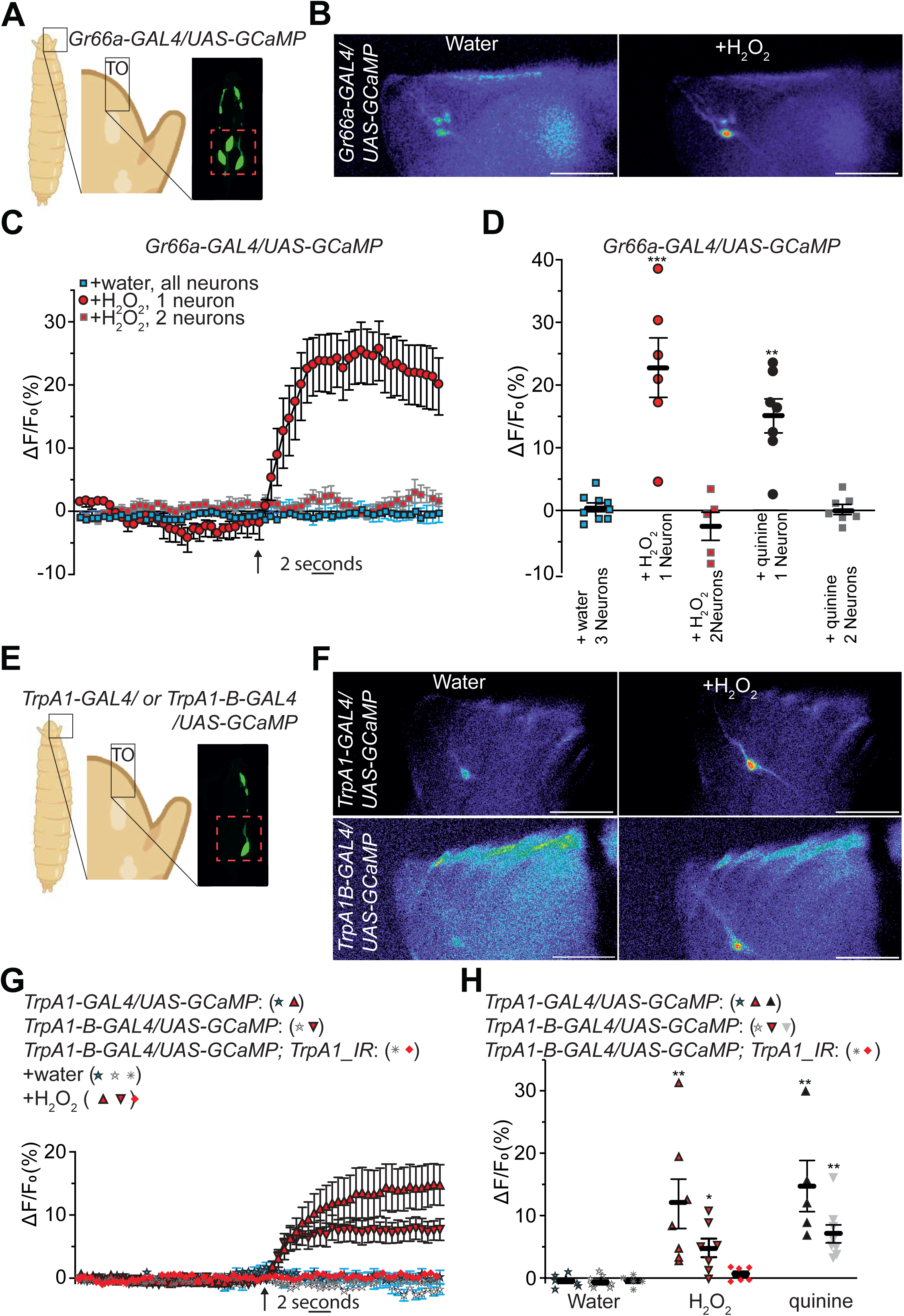
The *TrpA1+* neuron at the tip of the larval anterior part responds to H_2_O_2_ in a *TrpA1*- dependent manner. (**A** and **E**) Graphical representation of larval region observed during the GCaMP assay with 3 neurons detectable without stimulation in *Gr66a-GAL4/UAS-GCaMP7s* animals (**A**-**D**) and 1 neuron detectable without stimulation in *TrpA1-GAL4/UAS-GCaMP7s* larvae as well as in *TrpA1-B-GAL4/UAS-GCaMP7s* animals (**E**-**H**). (**B**-**D**) Addition of H_2_O_2_ triggers a calcium concentration increase in one out of the three *Gr66a+* cells detectable in the anteriormost area of the larvae. (**B**) Representative images showing the GCaMP intensity before and after addition of either the control water or the H_2_O_2_ 1%. (**C**) Averaged ± SEM time course of the GCaMP intensity variations (ΔF/F0 %) for *Gr66a+* neurons. The addition of water (n = 9 flies) or H_2_O_2_ 1% (n = 6 flies) at a specific time is indicated by the arrow. (**D**) Averaged fluorescence intensity of negative peaks ± SEM in response to water (n = 9), H_2_O_2_ 1% (n = 6), or quinine (10 mM) (n = 7). (**E-H**) Addition of H_2_O_2_ triggers a calcium concentration increase in the unique *TrpA1+* cell detectable in the anteriormost area of the larvae. (**F**) Representative images showing the GCaMP intensity before and after addition of either the control water or the H_2_O_2_ 1%. (**G**) Averaged ± SEM time course of the GCaMP intensity variations (ΔF/F0 %) for *TrpA1+* as well as *TrpA1-B+* neurons and *TrpA1-B* cells expressing *UAS-TrpA1_IR* (*TrpA1-B-GAL4/UAS-GCaMP7s; TrpA1_IR*). The addition of water (n = 5-7 flies) or H_2_O_2_ 1% (n = 6-7 flies) at a specific time is indicated by the arrow. (**H**) Averaged fluorescence intensity of negative peaks ± SEM in response to water (n = 5-7 flies), H2O2 1% (n = 6-7 fies), or quinine (10 mM) (n = 5-8 flies). Scale bar is 20µm. * Indicates p=0.0132, ** indicates p≤0.0098, *** indicates p≤0.009, Kruskal– Wallis H test was used and the reference for a given assay is the same genotype exposed to water. Further details can be found in the detailed lines, conditions and, statistics for the figure section.

### Uracil induces a Duox-dependent signal activating *TrpA1-B+* neurons in naïve larvae

To validate our model that uracil released by bacteria activates intestinal Duox resulting in the production of ROS that are then detected by the anteriormost *TrpA1-B+* neuron, we tested whether larvae exposed to uracil could release a Duox-dependent signal capable of activating *TrpA1-B+* neurons in naïve individuals, *i.e.*, larvae that had not been exposed to uracil or any prior stimulating cue (*TrpA1-B-GAL4/UAS-GCaMP*). To do so, we exposed in liquid larvae that have been raised on antibiotic-containing food to 1mM uracil or water for 3 hours. The resulting conditioned medium was assayed for calcium responses in naïve, non–uracil-exposed animals (Figure 6A). Strikingly, while neither uracil alone nor water from larvae not exposed to uracil induced any response, the medium conditioned by uracil-treated larvae triggered a calcium rise in the anteriormost *TrpA1-B+* neuron (Figure 6B and 6C). Importantly, when *Duox* expression was specifically reduced in enterocytes of the uracil-treated larvae (*mex-Gal4/UAS-Duox_IR*), the conditioned medium no longer activated the *TrpA1-B+* neuron (Figure 6C). These results indicate that uracil exposure generates a Duox-dependent signal in donor larvae—likely a reactive oxygen species such as H₂O₂—that can activate *TrpA1-B+* neurons in naïve recipients.

**Fig. 6.**
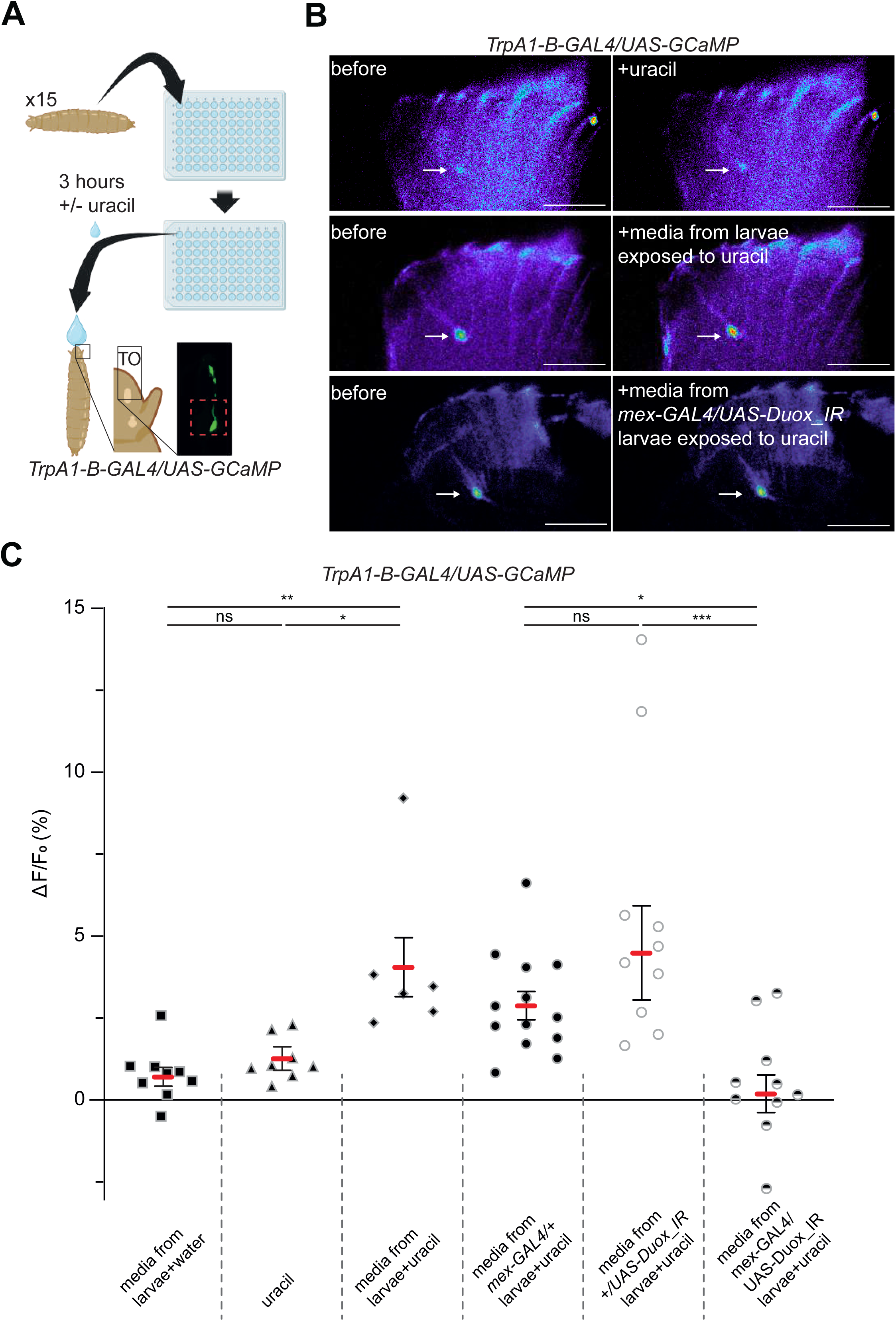
The *TrpA1-B+* neuron responds to medium conditioned by larvae exposed to uracil. (**A**) Graphical representation of the experimental design for the larva-conditioned medium. 15 larvae previously raised on antibiotic-containing medium are transferred at the L3 stage in a well containing either 100 µL of water or 100 µL of 1mM uracil diluted in water for 3 hours. After 3 hours in the dark at 25 °C, 50 µl of the medium in which the larvae were incubated is tested for its ability to trigger a calcium rise in the anterior part of *TrpA1-B-GAL4/UAS-GCaMP7s* larvae, which we consider naïve since they were raised on antibiotics and were not previously exposed to the larva-conditioned medium or to any of the chemicals tested. (**B**) Representative images showing the GCaMP intensity in the anteriormost *TrpA1-B+* neuron before and after addition of 50 µl of the medium: uracil alone or media form *w^-^* larvae exposed 3h to uracil or media form *mex-GAL4/UAS-Duox_IR* larvae exposed 3h to uracil. (**C**) Averaged fluorescence intensity of negative peaks ± SEM int the *TrpA1-B+* anteriormost neuron in response to either the chemicals used in the conditioned media assay (water, uracil) or the medium in which the larvae were incubated in (media form *w^-^* larvae+water, media form *w^-^* larvae+uracil 1mM, media form *mex-GAL4/+* larvae+uracil 1mM, media form *+/UAS- Duox_IR*+uracil 1mM, media form *mex-GAL4/UAS-Duox_IR*+uracil 1mM). Scale bar is 20µm. *p < 0.05, **p ≤ 0.0012, ***p ≤ 0.0002. Statistical analyses were performed using the Kruskal– Wallis test for assays involving *w^-^* animals (non-normal distributions) and one-way ANOVA followed by Tukey’s multiple-comparisons test for assays involving GAL4/UAS lines (after confirmation of normality). Further details can be found in the detailed lines, conditions and, statistics for the figure section.

### Free-feeding assays confirm PGN avoidance and reveal its dependence on larval microbial exposure and TrpA1

Using the PER method, we showed that adult avoidance of PGN relies on larval exposure to bacteria that produce uracil and is mediated by ROS and specific TrpA1 isoforms in host *Gr66a+*neurons. However, since proboscis extension reflects only one stage of a complex feeding sequence involving smell, tarsal detection, ingestion, and pharyngeal evaluation, we asked whether this aversion also manifests in a more physiological and ecologically relevant context. We therefore turned to a two-choice free-feeding paradigm in which flies can move freely, explore the environment, and voluntarily ingest the substrate of their choice. Using a 96-well assay with colored agar [52], we assessed preference by scoring abdominal coloration after ingestion (Fig. 7A and B). Control (*w^-^*) females do not discriminate between food sources when 1mM sucrose is associated with a blue or red dye (preference index: P.I. = 0.5), flies with purple abdomen were observed (Figure 7B and 7C). However, animals robustly preferred 1 mM sucrose over a sucrose+PGN mixture, displaying a strong preference index for sucrose alone (P.I. >0.8) (Figure 7C). To determine whether larval microbial experience was required for the emergence of this adult PGN avoidance, we tested flies raised on antibiotics throughout the larval stage. These individuals showed a markedly reduced avoidance of sucrose+PGN (P.I. <0.55), with most animals ingesting both solutions. Finally, to evaluate the contribution of TrpA1—as anticipated from our PER results—we tested *TrpA1-KO* flies. These mutants consumed sucrose and sucrose+PGN equally, resulting in a strongly diminished preference index toward sucrose alone (P.I.=0.51) (Figure 7C). Finally, the *TrpA1-BCE-KO* animals did not distinguish sucrose alone from the sucrose mixed with PGN (P.I.=0.51) while *AD-KO* did make a discrimination, indicating that *-A* and *-D* isoforms are dispensable and suggesting *-B* and/or *-C* and/or *-E* as necessary.

**Fig. 7.**
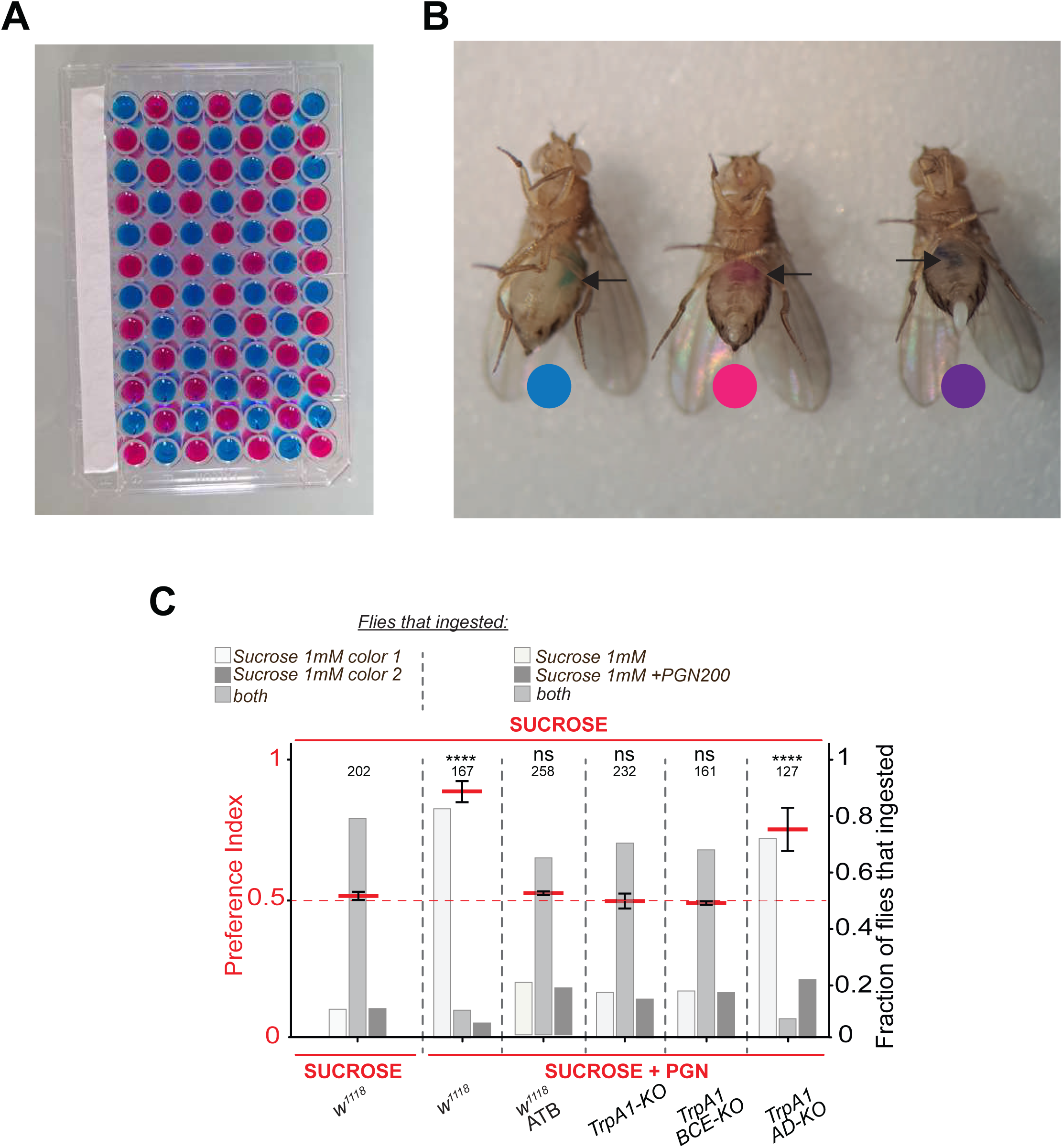
PGN is avoided by free-moving animals in a 2-choices assay. (**A-C**) Larval cohabitation with bacteria and the TrpA1 channel are necessary for a PGN- triggered aversion in a 2-choices assay. (**A**) The plates with the dye and (**B**) the flies that ingested the colored media leading to (**C**) the preference indexes for different conditions and genotypes as well as the fraction of flies that ingested the proposed mixtures. For (**C**), the averaged preference index is in red with the standard error of the mean obtained from 6 to 8 independent experiments. The total amount of females tested is indicated on top of each assay. A preference index (PI) over 0.5 indicates a preference toward sucrose while a PI of 0.5 indicates either that two opposite behaviors occurred in the population or that a majority of animals consumed both solutions. The fraction of flies that ingested one, the other or both solutions allow to discriminate. In light colors are the fractions of flies that ingested one solution or both solutions. The animals from all the experiments were pooled for statistical analyses. ns indicates p>0.05, **** indicates p<0.0001 Chi-square Test. Further details can be found in the material and methods section and in the source data file.

Together, these results demonstrate that adult female flies avoid ingesting sucrose+PGN mixtures even under free-feeding conditions and that this avoidance requires prior larval cohabitation with bacteria. Moreover, TrpA1 appears to be crucial in limiting PGN ingestion, probably due to its role in the sensory priming that occurs during larval microbial exposure.

## Discussion

Our previous work demonstrated that certain bacteria have the ability to *prime* Drosophila larvae, leading to the emergence of adults whose gustatory system perceives bacterial PGN as an aversive molecule [37]. The objective of the present study was twofold: first, to identify the bacterial molecules capable of inducing this phenomenon, and second, to characterize the host proteins and cells through which the signal is received and transmitted.

The observation that two bacterial strains differing only in their ability to secrete uracil exhibit distinct priming properties highlights the crucial role of uracil in this process. Interestingly, the same metabolite appears to participate in several mechanisms underlying interactions between pathogenic bacteria and their hosts [44], [45], [53], [54]. A more systematic analysis of the correlation between uracil production capacity and priming ability will help strengthen this finding and may reveal alternative priming mechanisms. Nonetheless, the fact that simple supplementation of the growth medium with uracil is sufficient to prime axenic larvae demonstrates that this bacterial signal is not only necessary but also sufficient to trigger the response. In this context, it will be important to determine the environmental factors that regulate and modulate bacterial uracil production.

Our results suggest that uracil acts through the ROS production system, with this step taking place in the digestive tract. Indeed, in the absence of Duox activity specifically in the gut, bacteria lose their ability to prime larvae. The observation that a similar blockage occurs upon overexpression of catalase under the *mex*-*GAL4* promoter supports the idea that ROS must be generated in the gut, and further suggests that hydrogen peroxide is the main ROS involved. Having identified ROS as the secondary messenger, we next asked which receptor and cell types are responsible for sensing and transmitting this signal. Using genetic tools enabling expression of specific TrpA1 isoforms, we identified a single neuronal cell located in the terminal organ of the anterior extremity, co-expressing the TrpA1-B isoform and the Gr66a protein, as essential for priming. Selective inactivation of *TrpA1* in this neuron completely abolishes priming by bacteria that otherwise induce it in wild-type larvae. Moreover, expression of the *TrpA1-B* isoform—but not *TrpA1-A*—in this neuron alone is sufficient to rescue the phenotype of a pan-*TrpA1* mutant, indicating that the different isoforms differ in their ability to respond to ROS, as previously shown for other ligands such as heat or citronella. Our data therefore suggest that bacterially derived uracil triggers Duox-dependent ROS production in the gut, which in turn activates TrpA1. Since *TrpA1* is also expressed in enteroendocrine cells, these were initially considered to be plausible mediators of ROS effects. However, our genetic analyses rule out this hypothesis and demonstrate that the *TrpA1+/Gr66a+* neuron in the terminal organ is required.

This raises the question of the mechanistic link between ROS produced in the gut and activation of the terminal organ neuron. One possibility is that ROS diffuse directly across the gut wall to reach the target neuron; however, this seems unlikely given the presence of enzymatic systems that rapidly degrade ROS. Alternatively, intermediate signaling steps could exist between intestinal ROS production and TrpA1 activation, implying that ROS are not the direct ligands of TrpA1 in the neuron. Our new results support a third, non–mutually exclusive mechanism in which ROS generated in the gut are released into the external environment and subsequently sensed by the anteriormost TrpA1-B+ neuron. We show that larvae exposed to uracil produce a Duox-dependent signal capable of activating TrpA1-B+ neurons in naïve individuals, as evidenced by calcium responses elicited by conditioned medium from uracil-treated larvae. Importantly, this activity is abolished when Duox is specifically knocked down in enterocytes of the donor larvae, indicating that intestinal Duox activity is required for generation of the signal. These findings suggest that a diffusible molecule—likely a reactive oxygen species such as H₂O₂—is released from the gut and can act at a distance to activate TrpA1-B+ neurons. Such a mechanism implies that gut-derived ROS can function as inter- individual cues, potentially allowing infected larvae to transmit information about microbial presence to nearby conspecifics and thereby influence population-level behavior.

## Acknowledgments

We thank Annelise Vialat-Lieutaud. This work was supported by CNRS, ANR Pepneuron (ANR-21-CE16-0027) to J.R. and Y.G. Research in Y.G.’s laboratory is supported by the CNRS, the “Université de Bourgogne Europe”, the Conseil Régional Bourgogne Franche-Comté (PARI grant), the FEDER (European Funding for Regional Economical Development), and the European Council (ERC starting grant, GliSFCo-311403).

## Materials and Methods

### STAR METHODS

#### KEY RESOURCES TABLE

**Drosophila strains**

w1118 (w−), BL#5905

*mex-GAL4*, BL#91367

*UAS-Duox-IR*, BL#38907

*mex-GAL4; Tub-GAL80ts* (*Tub-GAL80ts* BL#7016)

*UAS-Catalase*, BL#24621

*TrpA1-KO, BCE-KO, AD-KO, BC-KO, E-KO*

*TrpA1-A-KI, -B-KI, -C-KI, -D-KI, -E-KI*

*TrpA1-GAL4, -A-GAL4, -B-GAL4, -C-GAL4, -E-GAL4* (gifts from Xiang lab)

*UAS-GFP6x*, BL#52262

*TrpA1-B-GAL4; UAS-GFP6x* (this study)

*Gr66a-LexA; LexAop-mCherry* (gift from Zhang lab)

*UAS-TrpA1-IR*, BL#36780

*Gr39a.b-GAL4*, BL#57632

*Gr66a-GAL4*, BL#28801

*Tub-GAL80ts; UAS-TrpA1-IR* (this study)

*UAS-TrpA1-B; TrpA1−KO* (this study)

*UAS-TrpA1-A; TrpA1−KO* (this study)

*Gr66a-GAL4; TrpA1−KO* (this study)

*UAS-GCaMP7s* (gift from Matthieu Cavey)

*UAS-GCaMP7s; UAS-TrpA1-IR* (this study)

**Bacterial strains**

*Lactobacillus brevis* (gift from François Leulier’s lab)

*Erwinia carotovora* subsp. *carotovora* 15 (*Ecc15*)

*Ecc15 pyrE::Tn5*

*Ecc Δ4* (gifts from Won-Jae Lee lab)

**Chemicals**

Peptidoglycan (PGN-ECndi ultrapure), InvivoGen #tlrl-kipgn

Sucrose, Carl Roth #4621.1

Hydrogen peroxide, Sigma-Aldrich #H3410, #H1009

L-ascorbic acid (Vitamin C), Sigma-Aldrich #A92902

Uracil, Sigma-Aldrich #U0750

Quinine, Sigma-Aldrich #Q1125

**Software**

FIJI

GraphPad Prism 8 Leica MM AF 2.2.9

Adobe Photoshop BioRender

#### Resource Availability

##### Lead Contact

Further information and requests for resources and reagents should be directed to Julien Royet and will be fulfilled by the Lead Contact.

##### Materials Availability

All fly lines generated in this study are available upon reasonable request.

##### Data and Code Availability

All data supporting the findings of this study are available in the Source Data files associated with the figures. No custom code was generated.

#### EXPERIMENTAL MODEL

##### Drosophila melanogaster

Flies were maintained at 25°C on a yeast/cornmeal medium under a 12 h light/dark cycle unless otherwise stated. Female flies aged 5–7 days were used for adult behavioral experiments. Germ-free and monoassociated animals were generated as described below.

##### Fly Stocks and Genetic Tools

All Drosophila melanogaster strains used in this study are listed in the Key Resources Table. Unless otherwise stated, w1118 (w−) flies were used as the reference genetic background. *GAL4/UAS*-based genetic manipulations were performed using standard crosses. Newly generated fly lines were verified by genotyping and/or functional validation. All *TrpA1* mutant, knock-in, and *GAL4* lines were obtained from the Xiang laboratory and maintained according to their recommendations.

##### Fly Rearing Conditions and Diet Composition

Flies were reared at 25°C in temperature- and humidity-controlled incubators under a 12 h light/12 h dark cycle. Standard yeast/cornmeal food was prepared by boiling agar (8.2 g/L), cornmeal flour (80 g/L), and yeast extract (80 g/L) for 10 min. After cooling, methylparaben sodium salt (5.2 g/L) and propionic acid (4 mL/L) were added. This diet is protein-rich and sugar-poor, allowing the growth of a complex microbial community. For germ-free conditions, antibiotics (kanamycin, tetracycline, ampicillin, erythromycin) were added to the medium. For chemical supplementation experiments, 200 µL of Vitamin C (0.2 mg/mL) or hydrogen peroxide (1%) was added directly onto eggs laid for 6 h.

##### Bacterial Stocks and Culture Conditions

*Ecc15* strains were grown on LB agar plates at 30°C for at least 18 h. *L. brevis* was cultured on MRS agar at 37°C for at least 48 h under anaerobic-like conditions. Liquid cultures were initiated from single colonies and grown overnight (*Ecc15* in LB with shaking at 30°C; *L. brevis* in MRS broth, static, 37°C). Cultures were centrifuged at 4200 rpm for 15 min and adjusted to OD600 = 1. Heat-killed bacteria were prepared by incubation at 95°C for 5 min, and killing efficiency was verified by plating. Supernatants were obtained by centrifugation followed by filtration through a 0.45 µm filter.

##### Generation of Germ-Free and Monoassociated Flies

Embryos were collected from apple agar plates and sterilized using sequential washes in bleach (2.6%), ethanol (70%), and sterile water. Sterile embryos were transferred onto autoclaved food in sterile Falcon tubes. Monoassociation was achieved by adding 200 µL of bacterial suspension, supernatant, or control broth onto sterile food.

##### RNAi and Stage-Specific Genetic Manipulations

RNAi experiments were conducted using F1 progeny from *GAL4* × *UAS-RNAi* crosses. For temporal control of gene expression, *Tub-GAL80ts* was used. *GAL4* activity was suppressed at 18°C and induced at 29°C. For larval-specific RNAi, animals were shifted to 18°C at early pupal stages; for adult-specific RNAi, animals were shifted to 29°C after eclosion.

##### Proboscis Extension Reflex (PER) Assay

Female flies aged 5–7 days were starved for 24 h with access to water. Flies were anesthetized on ice, immobilized on adhesive tape, and allowed to recover for 1.5 h in a humid chamber. PER was elicited by contacting the labellum with filter paper soaked in control or test solutions following a defined stimulation sequence. Flies failing to respond to control sucrose stimulations were excluded from analysis.

##### Aversion and Two-Choice Feeding Assays

Aversion assays were performed using 1 mM sucrose as control and PGN-containing solutions as test stimuli. Two-choice feeding assays were conducted using dye-labeled agarose wells, and preference indices were calculated based on abdominal coloration.

##### Larval-Conditioned Medium Preparation

Conditioned media were generated by incubating 15 larvae in water or uracil (1 mM) for 3 h. Media were collected and immediately used for calcium imaging assays.

##### Calcium Imaging

Early third instar larvae expressing *GCaMP7s* were immobilized between coverslips. Fluorescence imaging was performed using a Lumencor light source and Hamamatsu ORCA- Flash 4.0 camera. Images were acquired every 500 ms, and ΔF/F was calculated after background subtraction using FIJI.

##### Microscopy

Whole larvae or dissected tissues were mounted in Vectashield and imaged using a Zeiss LSM 780 confocal microscope with a 20× air objective.

##### Quantification and Statistical Analysis

Statistical analyses were performed using GraphPad Prism 8. Normality was assessed using the D’Agostino–Pearson test. Non-parametric tests (Kruskal–Wallis) were applied where appropriate. PER data were analyzed using Fisher’s exact test with 95% confidence intervals. At least three independent biological replicates were performed for behavioral assays.

#### METHODS DETAILS

##### Fly stocks

The reference strain in this study corresponds to *w1118* (*w^-^*) BL#5905, *mex-GAL4* BL#91367, *UAS-Duox_IR* BL#38907, *mex-GAL4; TubG80ts* (this study and [55]; *TubG80ts* is BL#7016), *UAS-Catalase* BL#24621, *TrpA1-KO* / *BCE-KO* / *AD-KO* / *BC-KO* / *E-KO* / *A-KI* / *B-KI* / *C-KI* / *D-KI* / *E-KI*, *TrpA1-GAL4* / *A-GAL4* / *B-GAL4* / *C-GAL4* / *E-GAL4* (all TrpA1 genetic tools were very kind gift from the Xiang lab with a lot of support and advices ([46]), *UAS-GFP6x* BL#52262, *TrpA1-B-GAL4, UAS-GFP6x* (this study), *Gr66a-LexA ; LexAop-mCherry* (gift from Zhang lab, [56]), *UAS-TrpA1-IR* BL#36780, *Gr39a.b-GAL4* BL#57632, *Gr66a-GAL4* BL#28801, *TubG80ts; UAS-TrpA1_IR* (this study), *UAS-TrpA1-B; TrpA1^-^* (this study), *UAS-TrpA1-A; TrpA1^-^* (this study), *Gr66a-GAL4; TrpA1^-^* (this study), *UAS-GCaMP7s* (Gift from Matthieu Cavey), *UAS-GCaMP7s; UAS-TrpA1_IR* (this study).

##### Fly culture

Flies were grown at 25 °C on a yeast/cornmeal medium in 12 h/12 h light/dark cycle-controlled incubators. For 1 L of food, 8.2 g of agar (VWR, cat. #20768.361), 80 g of cornmeal flour (Westhove, Farigel maize H1), and 80 g of yeast extract (VWR, cat. #24979.413) were cooked for 10 min in boiling water. 5.2 g of Methylparaben sodium salt (MERCK, cat. #106756) and 4 mL of 99% propionic acid (CARLOERBA, cat. #409553) were added when the food had cooled down. It is important to mention that our conventional media allows the presence of several bacterial species [37] and is a protein-rich and sugar-poor culture medium. For germ-free condition, we raised flies on antibiotic (ATB) media. A mix of four ATB is added to the conventional media: Kanamycin + Tetracycline + Ampicillin + Erythromycin. For the experiments with Vitamin C and H_2_O_2_, we add 200 µL of Vitamin C at 0.2 mg /mL on eggs laid for 6 hours on conventional media and 200 µL of H_2_O_2_ 1% on eggs laid for 6 hours on ATB media. We use a protein-rich rearing media, in contrast to the sugar-rich media used in some other laboratories. Consequently, our animals may be more sensitive to low sucrose concentrations (1 mM) compared to flies raised on sugar-rich media. Therefore, assays using 1 mM sucrose may not yield reproducible results when performed with flies reared on sugar-rich media.

##### Bacterial stocks

*Levilactobacillus brevis* (*L. brevis*) is a gift form François Leulier’s Lab.

Ecc^15^ [57] is *Erwinia carotovora* subsp. *carotovora 15* and *Ecc^15^ pyrE::tn5* [45]*, Ecc^15^ Δ4* [58]. Gifts from Won-Jae Lee Lab.

##### Bacterial culture

*Ecc^15^* was grown on standard LB agar plates at 30°C at least 18 hours and *L. brevis* was grown in MRS agar in anaerobic-like conditions at 37°C for at least 48 hours. A single colony was used to prepare liquid cultures. *E*cc^15^ was grown in 200mL LB media (Lennox, Sigma-Aldrich ref L3022 or L2897) at 30°C with agitation and *L. brevis* was grown in 50 mL MRS liquid media (Sigma-Aldrich ref 69964 agar and 69966 broth) static at 37 °C in sealed 50 mL tubes for anaerobic conditions. After overnight growth, the cultures were centrifuged for 15 min at 4200 rpm. Bacterial doses were adjusted by measuring culture turbidity at an optical density of 600 nm to set the culture to OD 1. For the heat kill experiment, we set the OD to 1 from an overnight incubated culture, then incubate this sample at 95°C for 5 minutes. We confirm the heat kill experiment using an MRS agar plate containing the sample before and after heat kill. For the supernatant experiment, we set the OD to 1 from an overnight-incubated culture, then centrifuged and the supernatant is then filtered with a 0.45 µm filter.

##### Proboscis Extension Reflex (PER) behavior

All flies used for the test were females between 5 and 7 days old. Unless experimental conditions require it, the flies are kept and staged at 25 °C to avoid any temperature changes once they are put into starvation. The day before, the tested flies are starved in an empty tube with water-soaked plug for 24 h at 25 °C. Eighteen flies are tested in one assay, 6 flies are mounted on one slide and in pairs under each coverslip. To prepare the slide, three pieces of double-sided tape are regularly spaced on a slide. Two spacers are created on the sides of each piece of tape by shaping two thin cylinders of UHT paste. To avoid the use of carbon dioxide flies are anesthetized on ice. Under the microscope, two flies are stuck on their backs, side by side, on same piece of tape so that their wings adhere to the tape. A coverslip is then placed on top of the two flies and pressed onto the UHT paste, blocking their front legs and immobilizing them. Once all slides are prepared, they are transferred to a humid chamber and kept at 25 °C for 1.5 h to allow the flies to recover before the assay. Flies are tested in pairs, the test is carried out until completion on a pair of flies (under the same coverslip), and then move on to the next pair. Before the test, water is given to each pair of flies to ensure that the flies are not thirsty and do not respond with a PER to the water in which the solutions are prepared. Stimulation with the test solution is always preceded and followed by a control stimulation with a sweet solution, to assess the fly’s condition and its suitability for the test. During the test small strips of filter paper are soaked in the test solution and used to contact the fly’s labellum (three consecutive times per control and test phase). Contact with the fly’s proboscis should be as gentle as possible. Ideally the head should not move. A stronger touch may prevent the fly from responding to subsequent stimulation. Based on the protocol needed the test is done following the sequence and the timing in the table below.

**Table 1.**
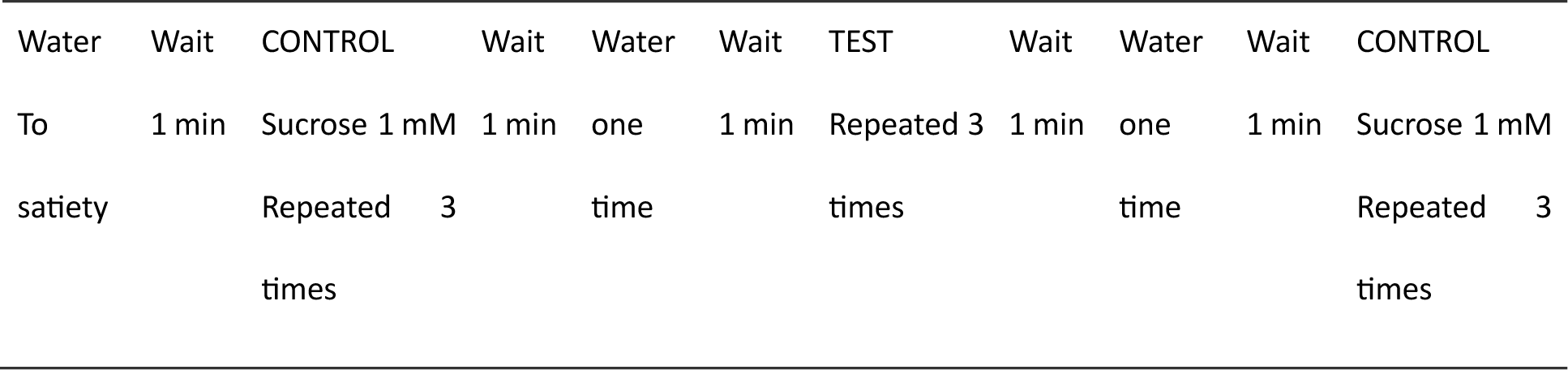
Aversion protocol.

All solutions to be tested are prepared the test day and stored at room temperature. In the aversion protocol, the control stimulation is performed with 1 mM sucrose (D(+)-sucrose ≥ 99.5%, p.a. Carl Roth GmbH + Co. KG). This concentration is sufficient to elicit a PER but is not so high as to influence the response to the subsequent test stimulation. PGN is *E. coli* K12: Invivogen, catalog code #tlrl-kipgn. After each control or test stimulation, a water-soaked strip is used to tap the proboscis and clean it. The response of the fly to each stimulation is recorded and averaged. We distinguish between two types of flies: those able to respond and those unable to respond. To make this distinction, we use three control stimulations before the test and three after the test. Flies that do not respond during the control stimulations (1 mM sucrose) are classified as “unable to respond” and are excluded from the analysis, regardless of their responses during the tests. Flies that respond during the control stimulations are classified as “able to respond,” and only their responses during the tests are included in the statistical analyses. The PER index is calculated as the percentage of flies tested that responded with a PER to the TEST stimulation and represented as ± 95% CI. In case of stage dependent experiments, flies are shifted from one condition to another upon hatching.

##### Monoassociation

An oviposition of *w*- germ free flies is set up on an apple agar plate with yeast at 25 °C for 6 hours. Three petri dishes are filled with 2.6% bleach, 70% ethanol (Ethanol 96° RPE Carlo Erba Ref 414638) and autoclaved purified distilled water respectively. The embryos are collected by filling the plate in which oviposition occurred with purified distilled water and using a small brush to gently detach them from the flies’ food. A 40 µm cell strainer is used to collect the embryos. The cell strainer with the embryos is then dipped into: bleach 2.6% for 5 min, ethanol 70% for 1 min, purified water for 1 min, ethanol 70% for 1 min, purified water for 1 min. The brush used to collect the embryos is sterilized in 2.6% bleach for ten min, rinsed and then used to transfer the sterile embryos onto the desired media.

We then deposit bleach eggs on ‘steril’ conventional media (medium is deposited in sterilized Falcon tubes in sterile conditions and sealed with a cap) and add 200µL of the monoassociation we want (MRS broth, MRS broth + *L. brevis* or supernatant or heat-killed bacteria, LB broth, LB broth + bacteria, LB broth + uracil).

##### RNAi

All the tested animals were F1 obtained from a cross between parents possessing the *GAL4* transgene and parents possessing the *UAS-RNAi* construction or from crosses between *w^-^* animals (genetic background of the *GAL4* and *UAS* lines) and a transgenic line to serve as ctrl. For stage-dependent RNAi experiments, the ubiquitously expressed *Tub-GAL80^ts^*, that inhibits the activity of *GAL4*, is temperature sensitive: it’s active at 18**°**C and inactivated at 29**°**C, allowing the expression of *UAS* when animals are raised at 29**°**C and preventing it at 18**°**C. For RNAi in larvae only, animals were raised from eggs to early pupae at 29**°**C and then shifted at 18**°**C. for RNAi in adults only, eggs, larvae and pupae were raised at 18**°**C and the virgin adults shifted to 29°C.

##### Larval-conditioned medium

Conditioned medium was used to highlight signals such as ROS produced by enterocytes in response to 1 mM uracil to potentially activate the *TrpA1-B+* neuron. In a 96-well plate (NUNC), conditioned medium was obtained by placing 15 larvae of adequate genotype, reared on antibiotic containing medium, in a well containing either 100 µL of water or 100 µL of 1mM uracil diluted in water for 3 hours. 50 µL of conditioned medium was then tested on the anterior part of *TrpA1-GAL4/UAS-GCaMP7s* larva reared on normal food in calcium imaging experiments. 100 µL of 1mM uracil was also placed in a well without any larvae during 3 hours to test the activity of 1mM uracil on the *TrpA1-B+* neuron localized in the terminal organ of *TrpA1-GAL4/UAS-GCaMP7s* larva.

##### Choice assay

Two-way choice assays were performed essentially as described [59]. Briefly, 5- to 7-day-old female flies (30 ± 10 flies per experiment) were starved for ∼18 h at 25°C and placed into a 96-well plate with 1% agarose in each well. Alternating wells contained either red (sulforhodamine B, 0.2 mg/mL; Sigma-Aldrich) or blue dye (bleu E133 meilleur du chef, 0.3%). Then we deposit 10µL of solution containing sucrose 1mM or Sucrose1mM + PGN 200 µg/ml on top of the well. Each experiment is carried out in duplicate by placing the components on two different colors (*e.g.*, sucrose 1mM on blue wells for plate A and on red wells for plate B) to verify whether there is an effect of color. The flies were allowed to feed in the dark for 90 min at ∼ 23°C. We transferred the plates to −20°C for 15min in order to rapidly freeze the flies in order to count the flies based on their abdominal color under a binocular microscope. The numbers of flies with blue, red, or purple (mixed red and blue) abdomens were tabulated (Fig 7A and 7B), and the P.I. was determined: (N_suc_ + 0.5N_mix_)/(N_suc_ + N_test_ + N_mix_). The flies without colors in their abdomen were discarded. The number of flies indicated corresponds to animals with colored abdomens.

##### Microscopy

No immunostaining was performed. To image larvae sensory organs and body as well as gut and brains, full larvae were killed with heat and mounted whole on slides using Vectashield fluorescent mounting medium. Organs were dissected in PBS, rinsed with PBS and directly mounted on slides using Vectashield fluorescent mounting medium. The tissues were visualized directly after. Images were captured with LSM 780 Zeiss confocal microscope (20x air objective was used). Images were processed using Adobe Photoshop and FiJi softwares.

##### Statistics

GraphPad Prism 8 software was used for statistical analyses. For in vivo calcium imaging, the D’Agostino–Pearson test to assay whether the values are distributed normally was applied. As not all the data sets were considered normal, non-parametric statistical analysis such as Kruskal–Wallis H test was used for all the data presented. For PER datasets. As the values obtained from one fly are categorical data with a *Yes* or *No* value, we used the Fisher exact t-test and the 95% confidence interval to test the statistical significance of a possible difference between a test sample and the related control. For PER assays, at least 3 independent experiments were performed. The results from all the experiments were gathered and the total amount of flies tested is indicated in the graph. In addition, we do not show the average response from one experiment representative of the different biological replicates, but an average from all the data generated during the independent experiments in one graph. However, each open circle represents the average PER of 1 experiment.

##### Chemicals

PGN-ECndi ultrapure peptidoglycan catalog code #tlrl-kipgn InvivoGen USA, D (+)-Saccharose ≥99.5 %.ref 4621.1 Carl Roth, H_2_0_2_ solution ref. H3410 (Figure 2 and 7) and H1009 (Figure S2) Sigma-Aldrich, Acid L-ascorbic (Vitamin C) ref. A92902 Sigma-Aldrich, Uracil ≥99.0 %. refU0750 Sigmal-Aldrich, Quinine ref. Q1125 Sigma-Aldrich.

##### Calcium imaging experiments

*In vivo* larval calcium imaging experiments were performed on beginning of third instar larvae. Larvae were immobilized in a small drop of distilled water placed between two plastic coverslips (22 mm × 22 mm, Agar Scientific). The two coverslips were held together to prevent movement of the anterior part of the larva using an alligator clip attached to a support. Stimulation was performed manually using a pipette with gel loading tip by applying water, Uracil (1mM) diluted in water, H_2_O_2_ (1%), quinine (10 mM, Sigma-Aldrich #Q11125) diluted in water or 50 µL conditioned medium through a hole made in the upper coverslip allowing the solutions to come into contact with the larval Terminal Organ (TO). For larval calcium imaging experiments, GCaMP7s was excited using a Lumencor diode light source at 482 nm ± 25. Emitted light was collected through a 505–530 nm band-pass filter. Images were collected every 500 ms using a Hamamatsu/HPF-ORCA Flash 4.0 camera and processed using Leica MM AF 2.2.9. Each experiment consisted of a recording of 70–100 images before stimulation and 160 images after stimulation. Data were analyzed as previously described [31] by using FIJI (https://fiji.sc/). For larvae fluorescence quantifications, a background fluorescence variation was calculated and subtracted to the fluorescence variation signal.

##### Detailed lines, conditions and statistics for the figure section

All these data are downloadable with the source data file.

Some schemes have been generated using Bio Render (https://www.biorender.com/).

## Competing Interests

The authors declare no competing interests.

## Declaration of generative AI and AI-assisted technologies in the writing process

During the preparation of this work the author(s) used Chat GPT to improve the readability and language of the manuscript. After using this tool/service, the author(s) reviewed and edited the content as needed and take(s) full responsibility for the content of the published article

## Supplementary figure legends

**Fig. S1.**
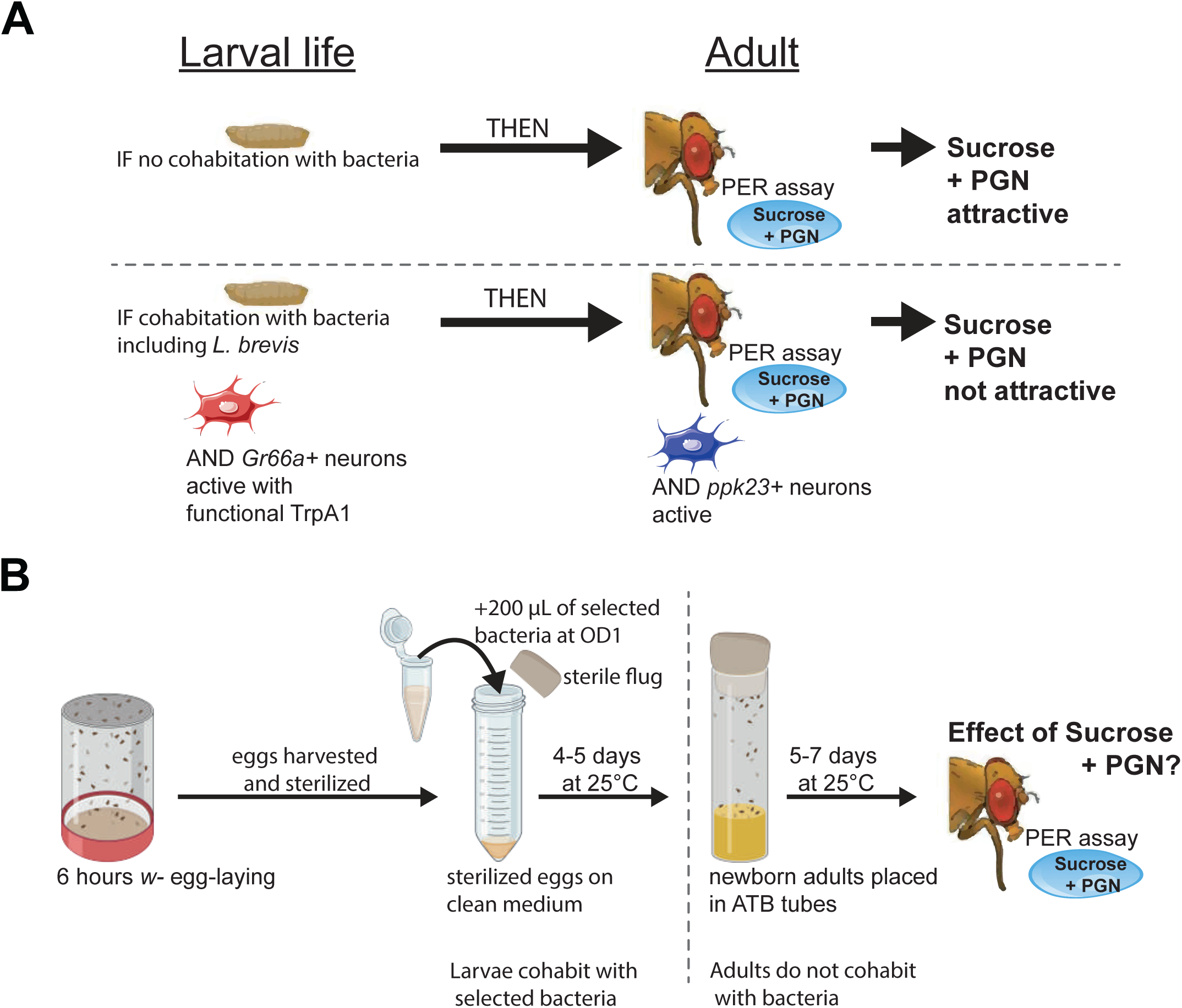
**(A)** Graphical representation of the priming model: adults derived from larvae raised without bacterial cohabitation do not reject sucrose + PGN mixtures. In contrast, adults from larvae cohabiting with bacteria in conventional media are attracted to sucrose but reject sucrose + PGN, a response dependent on larval *Gr66a+* neurons with functional *TrpA1* and adult *ppk23+* cells. **(B)** Graphical representation of the mono-association protocol used to expose larvae to a specific bacterial strain, followed by PER assay testing of the sucrose + PGN mixture in the resulting adults. Eggs are sterilized with bleach and deposited on a freshly made conventional media poured in a sterile culture tube and sealed with an autoclaved plug. Selected chemical such as LB or LB + uracil are added on top of the eggs, same with bacteria.

**Fig. S2.**
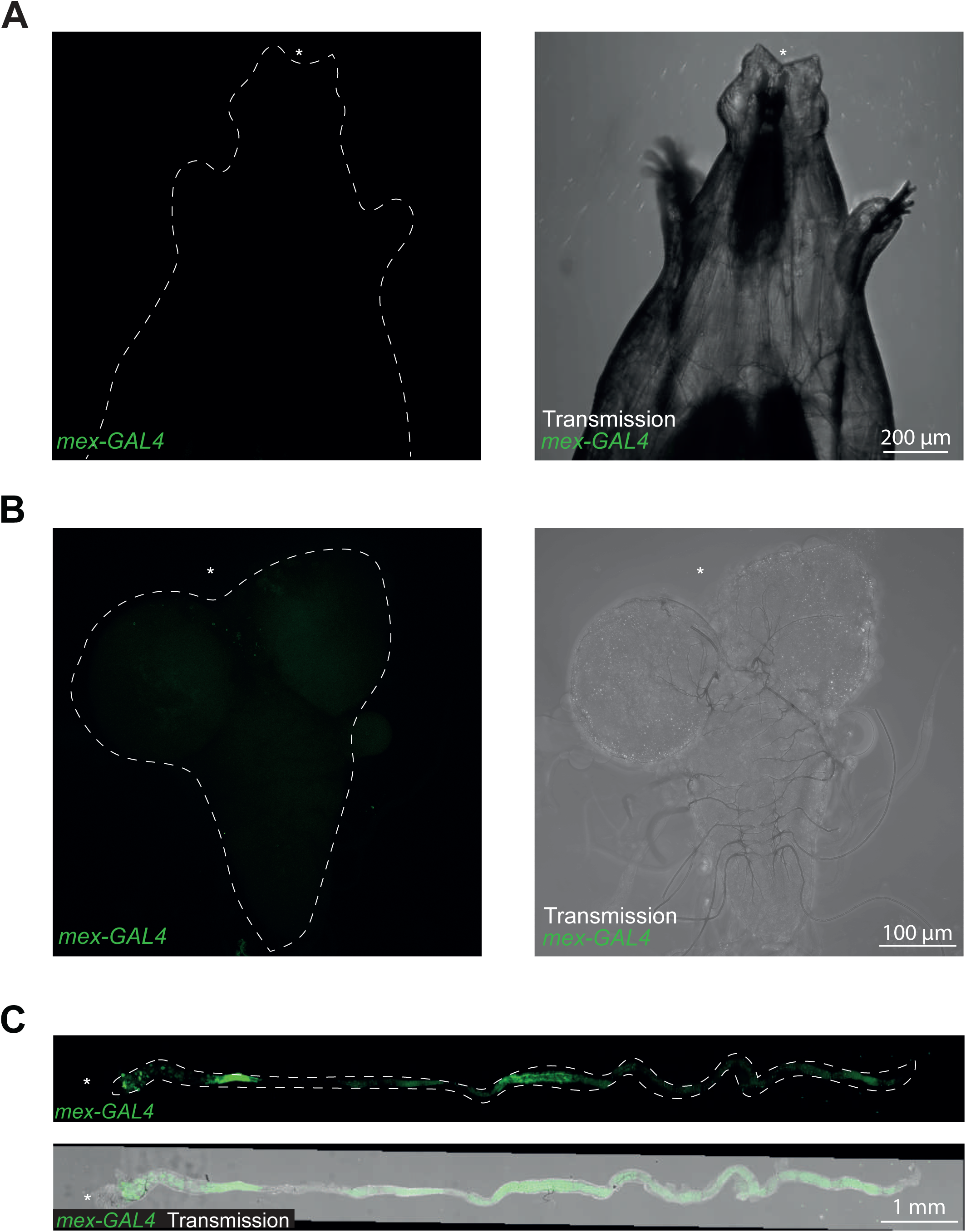
In larvae, the mex-*GAL4* driver is exclusively expressed in enterocytes. (**A-C**) Confocal images of larvae expressing *gfp* under the control of the *mex-GAL4* driver. Shown are representative pictures of the anterior extremities (**A**), the brains (**B**) and the guts (**C**). In (**A-C**), the anterior is indicated with an asterisk.

**Fig. S3.**
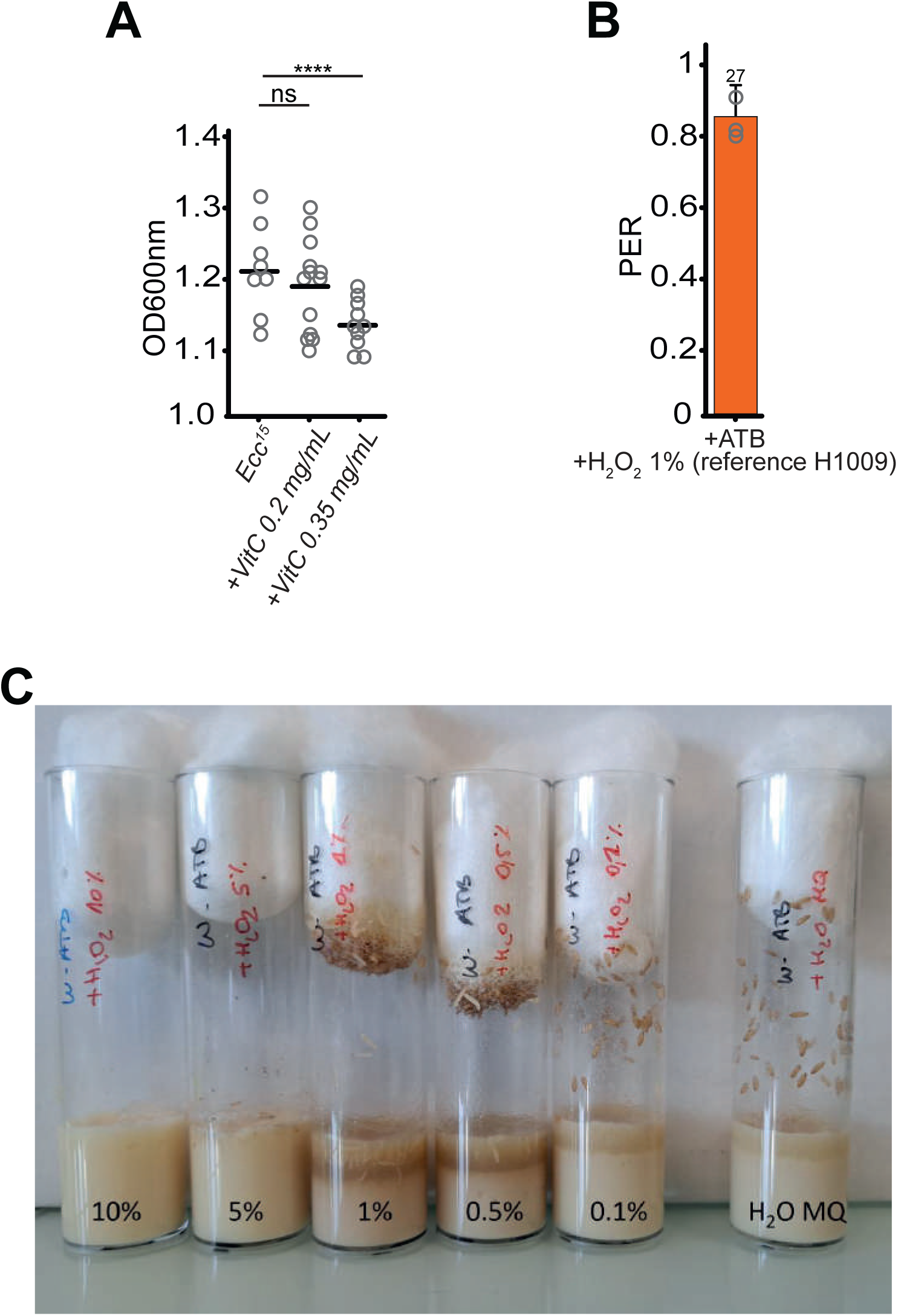
Vitamin C at 0.2 mg/ml is not bactericidal and H_2_O_2_ 1% does not impair larval development. (**A**) Vitamin C can be bactericidal. Vitamin C was added or not to a bacterial liquid culture and the OD 600nm was measured 24 hours later following an incubation at 30°C with agitation. ****p=0.056, non-parametric t-test, Mann-Whitney test (**B**) H_2_O_2_ supplementation with another chemical reference (H1009 contrary to H3410 used in Figure 2 and Figure 7) is not sufficient to prime. PER index to solutions of sucrose + PGN (PGN200) of *w^-^* flies. Larvae were raised on antibiotics-containing media with H_2_O_2_ (1%) using an H_2_O_2_ reference different form the one presented in Figure 2E. All the resulting adults were raised on antibiotics-containing media. The PER index is calculated as the percentage of flies tested that responded with a PER to the stimulation ± 95% confidence interval (CI). A PER value of 1 means that 100% of the tested flies extended their proboscis following contact with the mixture, a value of 0.2 means that 20% of the animals extended their proboscis. The number of tested flies (n) is indicated on top of the bar. At least 3 groups with a minimum of 10 flies per group were used. Each independent group is represented as an open circle. Further details can be found in the detailed lines, conditions and, statistics for the figure section. (**C**) H_2_O_2_ 1% (reference H3410) does not obviously impair larval growth. Images of tubes containing medium with antibiotics four days after bleached eggs deposition, followed immediately by addition of H_2_O_2_ at the indicated percentage.

**Fig. S4.**
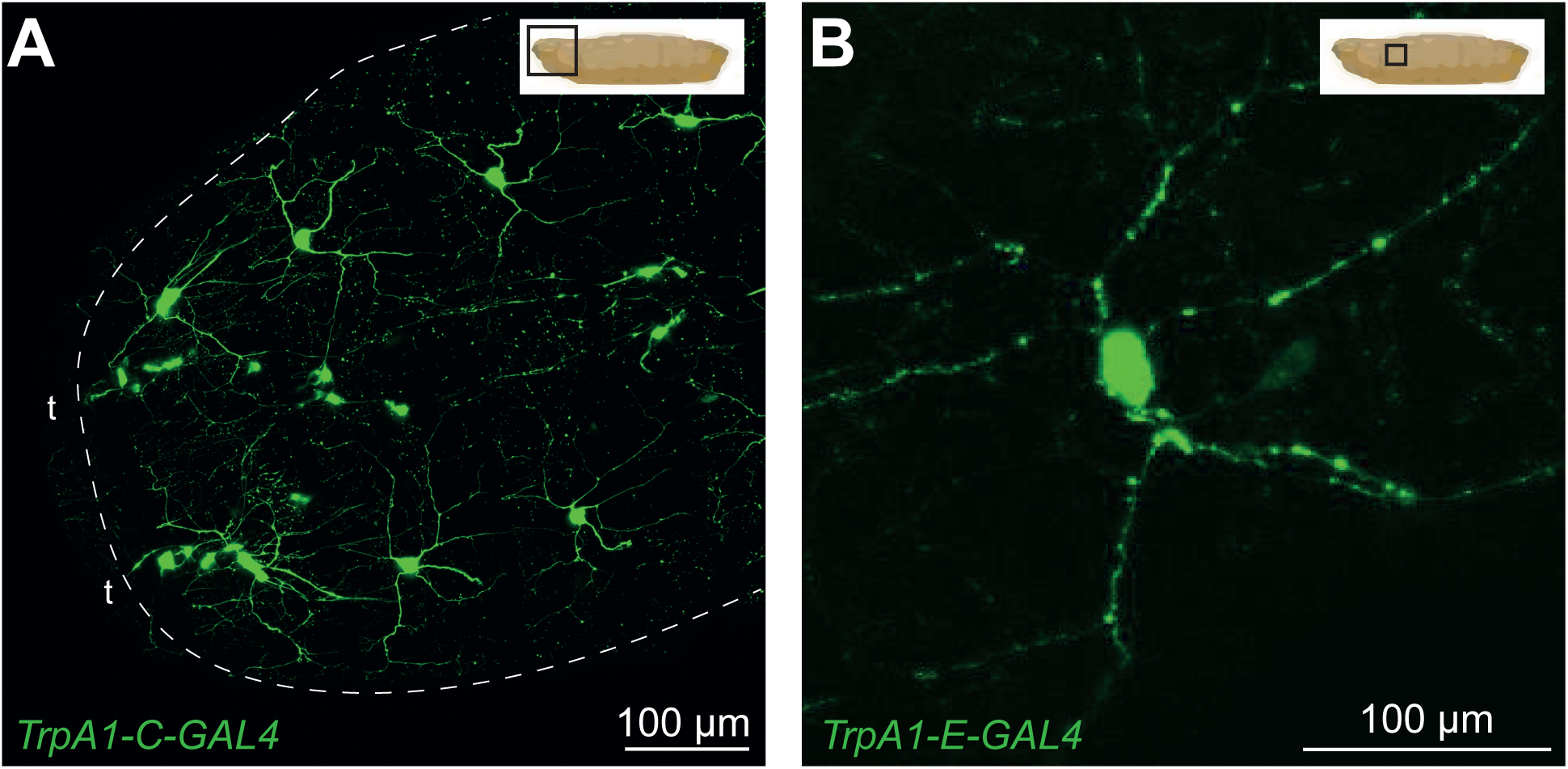
*TrpA1-C* and *TrpA1-E GAL4* drivers, but not *TrpA1-B-GAL4* are expressed in C4da neurons. (**A** and **B**) Representative confocal images of larvae expressing *gfp* under the control of the *TrpA1-C-GAL4* driver (**A**) or the *TrpA1-E-GAL4* driver (**B**). Shown are the C4da neurons in the anterior extremity with a dorsal view (**A**) or a magnified lateral view of a single C4da cell (**B**). In panels (**A**) and (**B**), anterior is oriented to the left, and a schematic representation of the whole larva indicates the depicted area. Neurons with dendrites extending toward the external environment are indicated by (t).

**Fig. S5.**
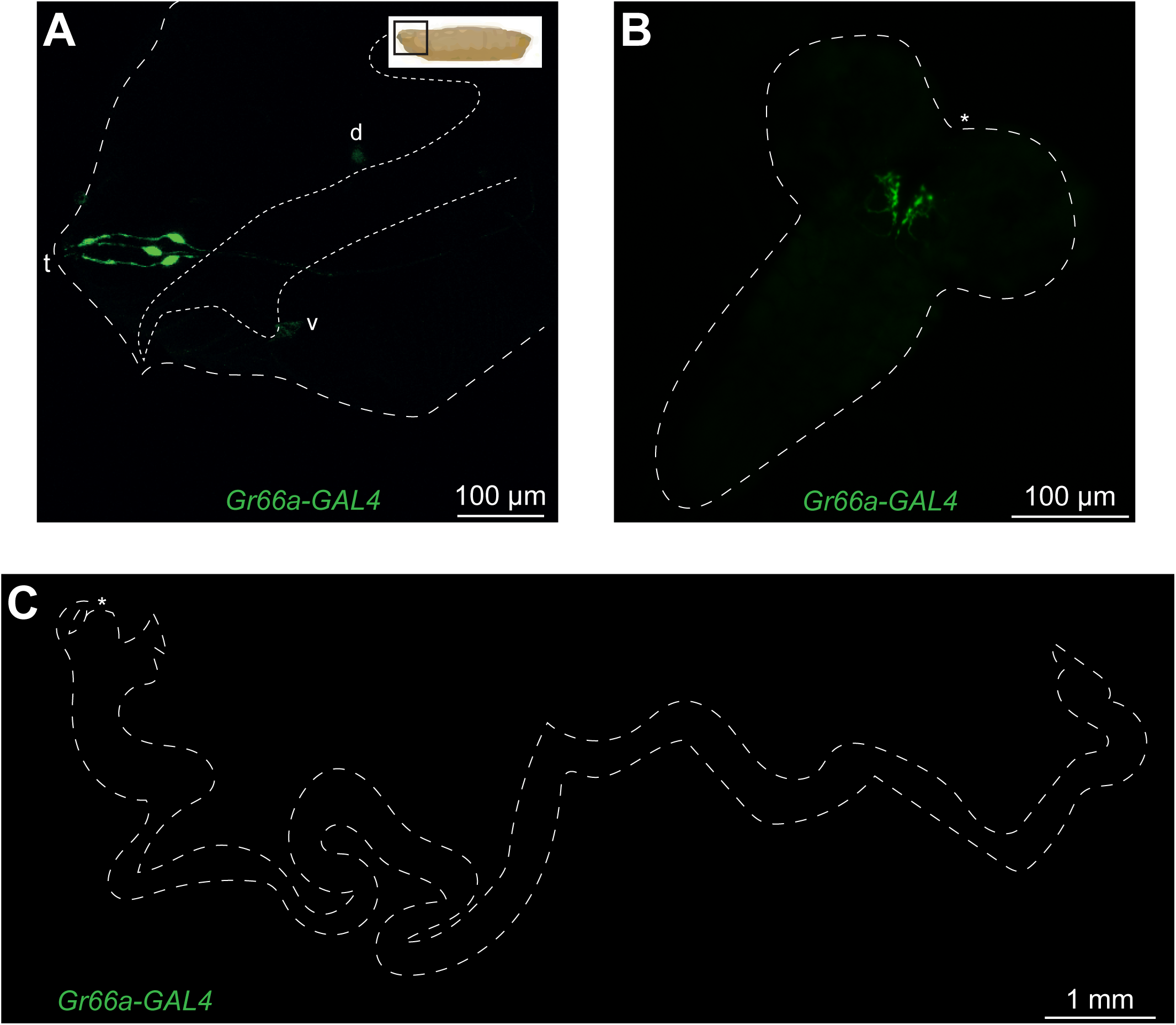
*Gr66a+* cells are present in larval anterior, brain, but not in the gut. (**A**-**C**) Representative confocal images of larvae expressing *gfp* under the control of the *Gr66a-GAL4* driver with the anterior extremity (**A**), the brain (**B**) and the gut (**C**). The anterior is either on the left (**A**) or indicated with an asterisk (**B** and **C**). Neurons with dendrites extending toward the external environment are indicated by (t) and despite being fainter dorsal pharyngeal sensilla ganglion (d) and ventral pharyngeal sensilla ganglion (v) are detectable. A schematic representation of the whole larva indicates the depicted area.

**Fig. S6.**
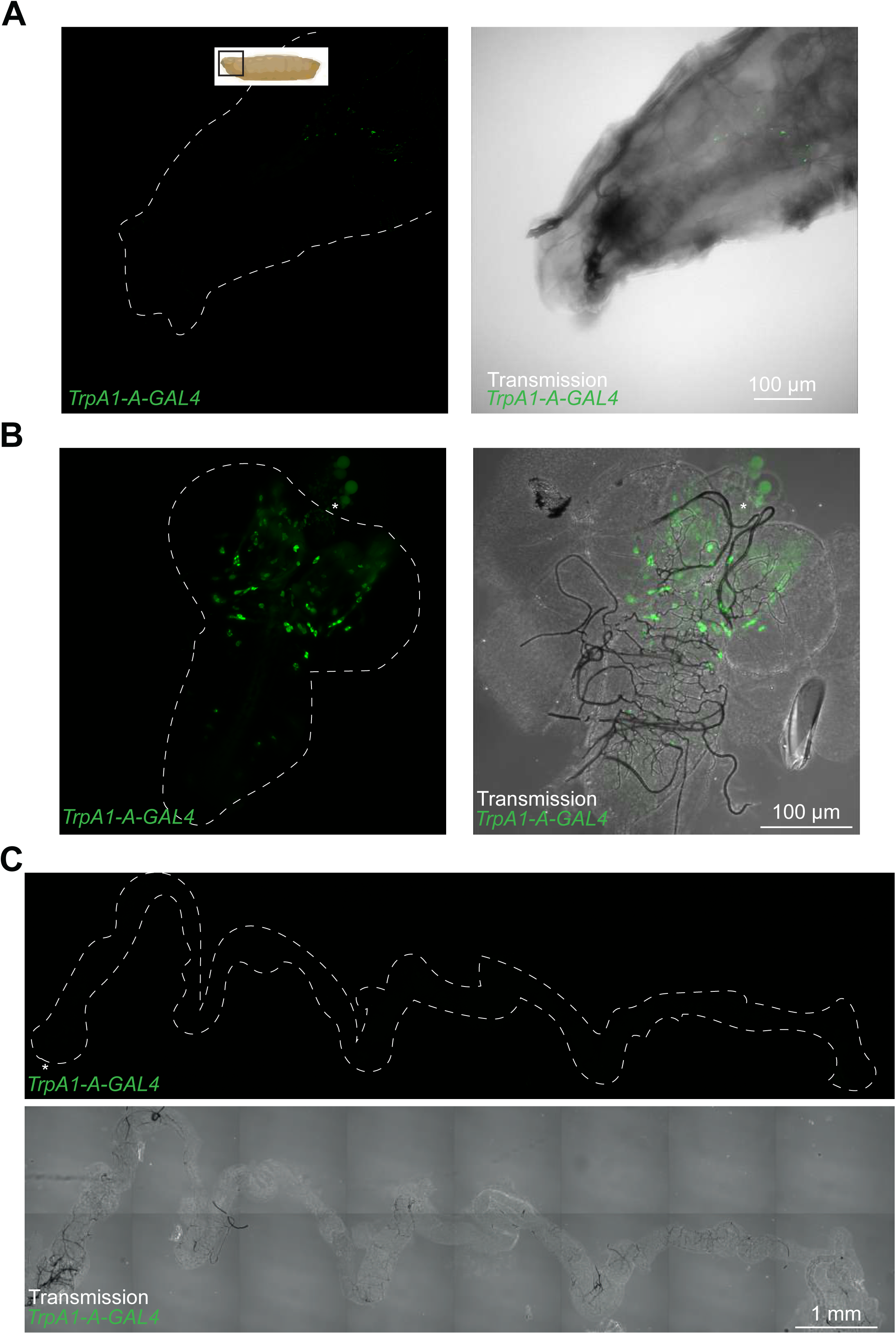
The *TrpA1-A* isoform is expressed according to an expression pattern that is different from that of the -*B*, -*C* and *-E* isoforms. (**A**-**C**) Representative confocal images of larvae expressing *gfp* under the control of the *TrpA1-A-GAL4* driver with the anterior extremity (**A**), the brain (**B**) and the gut (**C**). The anterior is either on the left (**A**) or indicated with an asterisk (**B** and **C**). A schematic representation of the whole larva indicates the depicted area.

**Fig. S7.**
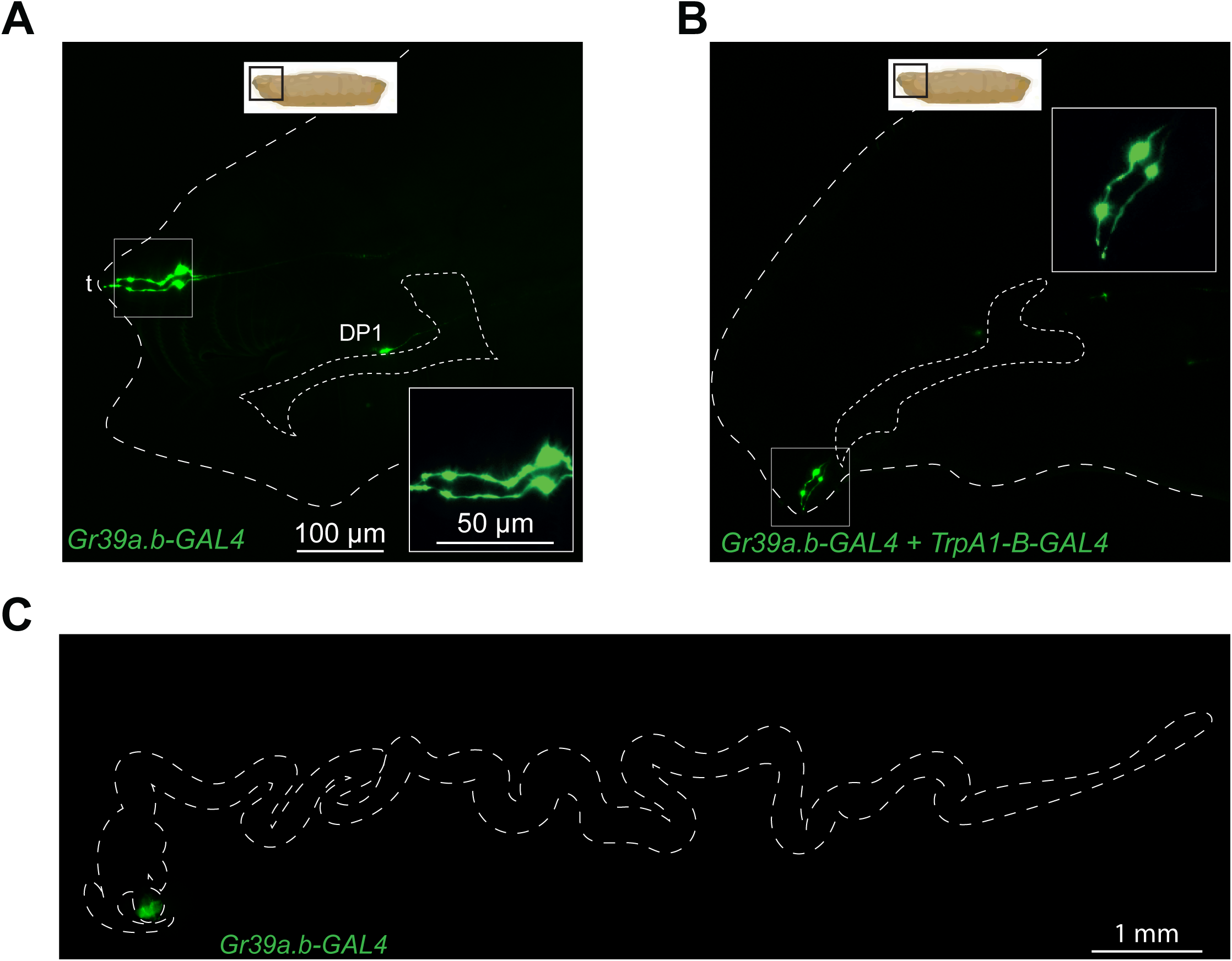
*Gr39a.b-GAL4* driver is only expressed in anterior neurons and includes TrpA1-B+ cell. (**A-C**) Confocal images of larvae expressing *gfp* under the control of the *Gr39a.b-GAL4* driver (**A**), the *TrpA1-B-GAL4* driver as well as the *Gr39a.b-GAL4* driver (**B**). Shown are the anterior parts (**A-B**) and the guts (**C**). In (**A-C**), the anterior is on the left and the larger square is a magnification of the most anterior part of the animal housing neurons whose dendrites extend toward the external environment (t), a schematic representation of the whole larva indicates the depicted area.

**Fig. S8.**
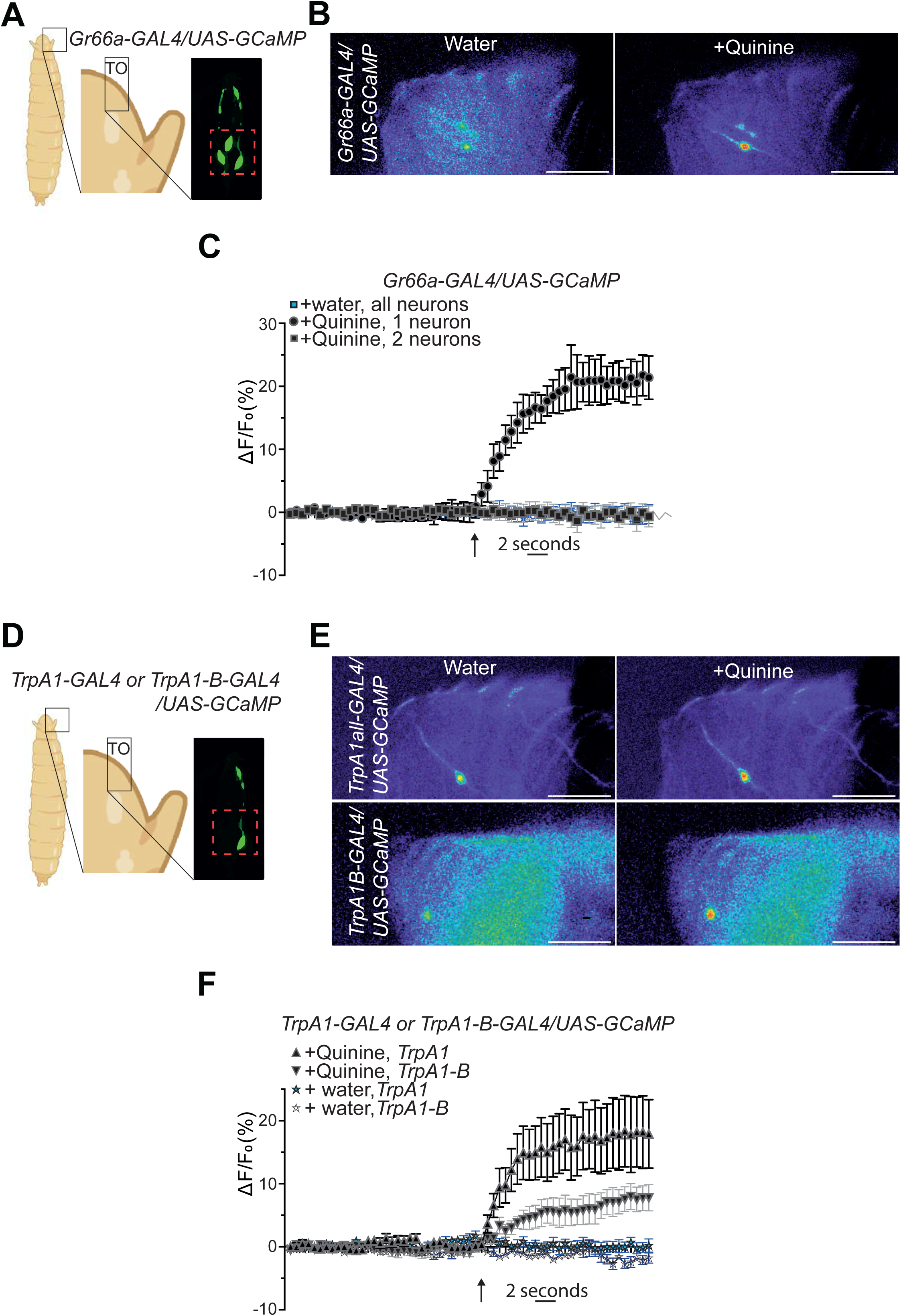
Quinine triggers a calcium increase in one *Gr66a+* neuron as well as in the anteriormost *TrpA1+* neuron. (**A** and **D**) Graphical representation of larval region observed during the GCaMP assay with 3 neurons detectable without stimulation in *Gr66a-GAL/UAS-GCaMP7s* animals (**A**-**C**) and 1 neuron detectable without stimulation in *TrpA1-GAL4/UAS-GCaMP7s* larvae as well as in *TrpA1-B-GAL4/UAS-GCaMP7s* animals (**D**-**F**). (**B** and **C**) Addition of quinine triggers a calcium concentration increase in one out of the three *Gr66a+* cells detectable in the anteriormost area of the larvae. (**B**) Representative images showing the GCaMP intensity before and after addition of either the control water or the quinine (10 mM). (**C**) Averaged ± SEM time course of the GCaMP intensity variations (ΔF/F0 %) for *Gr66a+* neurons. The addition of water (n = 9 flies) or quinine (n = 7 flies) at a specific time is indicated by the arrow. (**D**-**F**) Addition of quinine triggers a calcium concentration increase in the unique *TrpA1+* cell detectable in the anteriormost area of the larvae. (**E**) Representative images showing the GCaMP intensity before and after addition of either the control water or the quinine. (**F**) Averaged ± SEM time course of the GCaMP intensity variations (ΔF/F0 %) for *TrpA1+* as well as *TrpA1-B+* neurons. The addition of water (n = 5-8 flies) or quinine (n = 5-8 flies) at a specific time is indicated by the arrow. Scale bar is 20µm.

## Notes

### Competing Interest Statement

The authors have declared no competing interest.

